# Jumbo phage-mediated transduction of genomic islands

**DOI:** 10.1101/2024.08.29.610337

**Authors:** Yansong Zhao, Yue Ma, Christina Vasileiou, Andrew D. Farr, David W. Rogers, Paul B. Rainey

## Abstract

Bacteria acquire new genes by horizontal gene transfer (HGT), typically mediated by mobile genetic elements (MGEs). While plasmids, bacteriophages and certain integrative and conjugative elements (ICEs) are well characterized, the broader diversity of MGEs remains poorly understood. Here, we propagated the bacterium *Pseudomonas fluorescens* SBW25 in the presence of filtrate obtained from garden compost communities. Genome sequencing of derived colonies revealed acquisition of three different mobile elements, each integrated immediately downstream of *tmRNA*, flanked by direct repeats, and encoding a tyrosine integrase (*intY*) and putative phage defense systems. Absent are genes with recognized roles in autonomous transfer. Interrogation of DNA sequence databases showed that similar elements are widespread in the genus *Pseudomonas* and beyond, with *Vibrio* Pathogenicity Island-1 (VPI-1) from *V. cholerae* as a notable example. Bioinformatic analyses demonstrate frequent horizontal transfer among diverse hosts. Detailed analysis of a single element, I55, showed that it is transferred between cells by a jumbo phage, and confers a fitness benefit via a type II restriction-modification system.

**Significance Statement:** The impact of horizontal gene transfer on the evolution of bacteria outpaces that driven by spontaneous mutation, but knowledge of the range of mediators, the genes mobilized, and the mechanisms of movement have largely depended on inferences stemming from bioinformatics. Here we describe a real-time evolution experiment in which a single focal strain propagated with filtrate from a complex microbial community captured genetic elements carrying a diverse cargo of genes whose mobility was previously uncharacterized. The elements represent a widespread class of mobile DNA dependent only on a tyrosine integrase targeting a highly conserved genomic site. Genetic analysis of one element shows that it confers a significant fitness advantage via defense against bacteriophages, and hijacks a jumbo bacteriophage for intercellular transfer. Our findings reveal new insights into the hidden diversity and dynamics of MGEs in natural environments.

**Classification:** Biological Sciences / Microbiology

## Introduction

Mobile genetic elements (MGEs), such as bacteriophages, plasmids, and integrative and conjugative elements (ICEs), play a central role in shaping the structure and function of microbial communities (Arnold et al., 2022). In addition to driving their own replicative capacity, MGEs often capture and disseminate ecologically significant genes among bacterial hosts, facilitating horizontal gene transfer (HGT) (Arnold et al., 2022; Dmitrijeva et al., 2024; Quistad et al., 2020). Recent theoretical and experimental work suggests that accessory genes disseminated by MGEs may act as “community genes,” promoting feedbacks that underpin differential persistence and enabling selection to operate at the level of communities (Doolittle, 2024; Doolittle & Inkpen, 2018; Lenton et al., 2021; Rainey & Quistad, 2020; Roughgarden, 2020).

While classical genetics has led to discovery of various mechanisms of HGT, including conjugation (Lederberg & Tatum, 1946), transduction (Zinder & Lederberg, 1952) and transformation (Griffith, 1928), the majority of recent discoveries stem from genome analyses (Dmitrijeva et al., 2024; Groussin et al., 2021; Hackl et al., 2023). Comparisons among closely related strains frequently reveal regions with contrasting evolutionary histories, often termed “genomic islands” (GIs), which are inferred to have arisen via HGT (Koutsovoulos et al., 2022; Yuan et al., 2023). While instructive, such bioinformatic methods rarely provide insight into the causes of horizontal movement. Notable exceptions include demonstration of the mobility of ICEs (Colombi et al., 2017; J. T. Sullivan & Ronson, 1998), of plasmids (Hall et al., 2016; Lilley & Bailey, 1997b), of gene transfer agents (Lang et al., 2012; Marrs, 1974), and of phage-inducible chromosomal islands (PICIs) (Barcia-Cruz et al., 2024; Fillol-Salom et al., 2019; Lindsay et al., 1998; Novick et al., 2010). PICIs, a class of phage satellite (Ibarra-Chávez et al., 2021; Penadés et al., 2025), hitchhike with, or hijack, bacteriophages to facilitate their own intercellular transfer.

Experimental evolution, where populations or communities are propagated under controlled conditions (Lenski, 2017; Rainey et al., 2017; Rainey & Travisano, 1998), offers a powerful means to explore evolutionary processes. Although rarely used to study HGT directly, experimental evolution provides opportunities to track the dissemination of genes, identify the mediators of transfer, and uncover functional consequences (Frazão et al., 2019; Hall et al., 2016; Lilley & Bailey, 1997a; Quistad et al., 2020; Woods et al., 2020). For example, Quistad et al. (Quistad et al., 2020), used experimental evolution of compost-derived microbial communities to show that manipulation of MGE activity influenced the flux of horizontally disseminated DNA (van Dijk et al., 2023), but in the absence of culturable hosts carrying newly acquired DNA – underpinned by further analyses of fitness and genetics – understanding is limited to inferences stemming from bioinformatics.

In this study, we use experimental evolution to detect, isolate and characterize MGEs from environmental sources. Building on the approach of Quistad et al. (Quistad et al., 2020) we simplified the system by propagating a single bacterium, *Pseudomonas fluorescens* SBW25 (Fortmann-Grote et al., 2023; Silby et al., 2009), in the presence of filtrate derived from complex compost communities. This led to capture of three GIs lacking any evidence of autonomous capacity for horizontal transfer. Each element integrates immediately downstream of *tmRNA*, varies in size and gene content, and encodes putative phage defense systems. Bioinformatic analyses based on a minimal set of core genes revealed these elements to be widespread across *Pseudomonas* and beyond, with frequent signatures of horizontal transfer. Detailed analysis of a single element, I55, showed that it is transferred between cells by a jumbo phage present in the compost filtrate, and confers a fitness benefit via a type II restriction-modification system.

## Results

### *Pseudomonas fluorescens* SBW25 propagated in the presence of compost filtrate acquires mobile genetic elements

To investigate whether *P. fluorescens* SBW25 could acquire MGEs from environmental sources, we used a simple mark-recapture approach (Fig. 1a(ii)). Initially, SBW25 was marked with gentamicin resistance (strain MPB28806) and propagated under static conditions in microcosms containing a wash from compost communities. Unfortunately, SBW25 rapidly went extinct (Fig. E1). To reduce competition and risk of competitive exclusion, we propagated SBW25 carrying kanamycin resistance and either GFP or RFP markers (strains MPB29284 and MPB29300, respectively) in the presence of filtrate from compost communities, employing two regimes: a "live-filtrate" regime and a "fixed-filtrate" regime (Fig. 1a(i) and 1a(iii), respectively). In both cases, filtrate was prepared by passing community cultures through 0.2 μm filters to remove cells while retaining phage-sized particles and other small entities (Quistad et al., 2020).

**Fig. 1.**
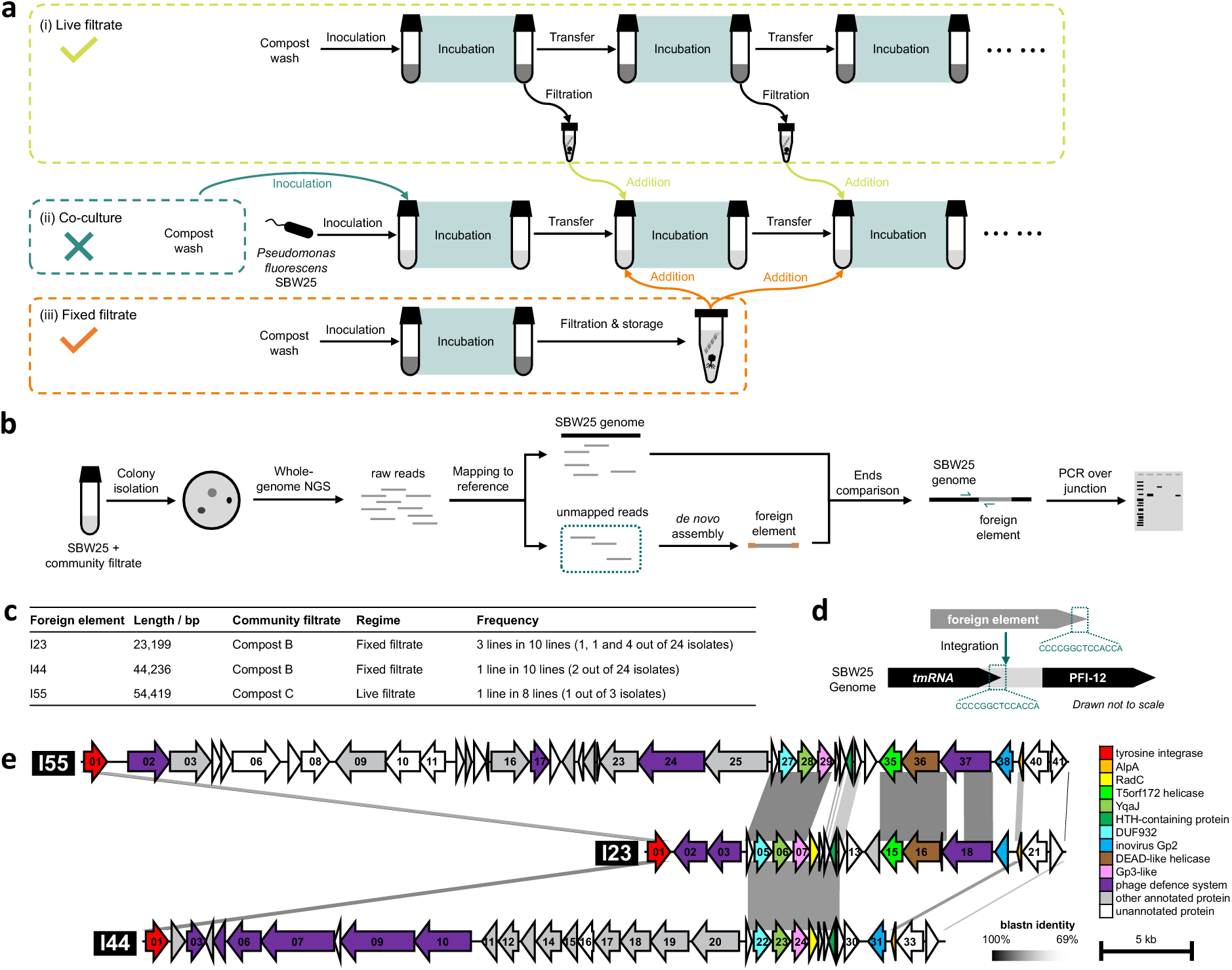
Experimental evolution and detection of foreign elements. **a.** Three serial-propagation regimes to evolve SBW25 populations. (i) live filtrate regime; (ii) co-culture regime; (iii) fixed filtrate regime. Components within each box are specific to each regime, and components outside of boxes are common to all regimes. **b.** The pipeline for screening and confirming SBW25 strains carrying foreign elements inserted into the genome. **c.** Basic information for the 3 foreign elements detected. The table lists the number of base pairs of each foreign element, the name of the compost community from which the community filtrate was generated, the serial-propagation regime, the number of parallel lines from which isolates with the foreign element were detected, and the number of isolates containing the foreign element in each such line. **d.** Insertion site of the foreign elements in the SBW25 genome. Upon insertion, two 14 bp direct repeats flank the foreign element. The direct repeats at the 5’-end and 3’-end of the element were treated as from SBW25 genome and from the foreign element, respectively, despite the exact origin of each repeat is unknown. **e.** The aligned genetic maps of the 3 foreign elements, I23, I44 and I55. The alignment was created by Easyfig (M. J. Sullivan et al., 2011). Parallelograms between MGEs indicate similar genomic regions based on blastn. The ORFs were numbered serially from 5-end to 3’-end for each MGE, and colored according to annotations. Detailed annotations of each ORF are listed in Table S4.

In the live-filtrate regime, material was freshly obtained from a compost community (compost C) synchronized with the evolution experiment, with 50 μL filtrate added to each of eight replicate SBW25 populations at every transfer over 14 serial passages. In the fixed-filtrate regime, filtrate was prepared once from independent compost communities (composts B, C, and D), stored at −80 °C, and added to 10 replicate populations per compost source for seven serial transfers. Following the final transfer, colonies were isolated from each replicate for whole-genome sequencing: two to three colonies per replicate in the live-filtrate regime, and pools of 24 colonies per replicate in the fixed-filtrate regime.

Mapping colony (or pooled-colony) sequencing reads back to the SBW25 reference genome revealed the acquisition of foreign DNA sequences (Fig. 1b). Further analysis identified three distinct GIs, designated I23, I44, and I55 (approximately 23 kb, 44 kb, and 55 kb in size, respectively), recovered on five independent occasions (Fig. 1c). From the 19 colonies sequenced from the live-filtrate regime, one contained I55 (strain MPB33357), which was acquired from compost C. Of the 720 colonies sequenced from the fixed-filtrate regime, six contained I23 (strains MPB38392, MPB38393, MPB38394, MPB38395, MPB38396, MPB38895) and two contained I44 (strains MPB38906, MPB38907). I23 was acquired independently on three occasions from compost B; I44 was detected in just a single replicate receiving filtrate from compost B.

Mapping the ends of contigs containing each element confirmed that all three GIs inserted at the same genomic locus, immediately downstream of *tmRNA* and immediately upstream of a putative mobilizable genomic island PFI-12 (see Fig. 1d). This was confirmed by PCR analysis. Each element is flanked by a 14 bp direct repeat (CCC CGG CTC CAC CA), corresponding to the second half of the *attP* site described in *Pseudomonas aeruginosa* PAO1 (Fiedoruk et al., 2020). One copy of the 14 bp sequence is located in the SBW25 genome, spanning the final 11 bp of *tmRNA* and 3 bp immediately downstream, while the second copy is introduced upon integration of the mobile element.

Each element harbors a gene encoding a tyrosine integrase (IntY, I23-*orf01*, I44-*orf01*, I55-*orf01*) near the 5’-end and a gene encoding AlpA (I23-*orf20*, I44-*orf32*, I55-*orf39*), a putative transcriptional activator of *intY* (Barcia-Cruz et al., 2024; X. Wang et al., 2009), at the 3’-end (see Fig. 1e). It is likely that IntY catalyzes site-specific recombination of the MGE into the genome, with the 14 bp direct repeat being the common core sequence (Grindley et al., 2006; Groth & Calos, 2004). The integrases each show ∼20% divergence at the nucleotide level; *alpA* in I44 and I55 are identical, but ∼20% dissimilar to *alpA* from I23. Recombination thus impacts evolution of the elements.

Beyond the shared integration site and presence of *intY* plus *alpA*, the three elements share a ∼18 kb region of sequence similarity at the 3’-end, although I44 and I55 contain gaps (see Fig. 1e for alignment, and Table S4 for detailed annotations). The region encodes YqaJ (I23-*orf06*, I44-*orf23*, I55-*orf28*), an HTH-containing protein (I23-*orf11*, I44-*orf28*, I55-*orf32*), an inovirus Gp2-like protein (I23-*orf19*, I44-*orf31*, I55-*orf38*), along with many open reading frames predicted to encode proteins with no matches to either DNA or protein sequence databases. Some elements, notably I23 and I55, also encode a putative DNA methyltransferase predicted to function as a type II restriction-modification system (Tock & Dryden, 2005).

Between *intY* and the region of similarity – and accounting for length differences – are various cargo genes that differ among the three MGEs. However, all three elements encode genes predicted by DefenseFinder (Tesson et al., 2022) to confer protection against phages (Fig. 1e). I23 encodes a Gabija system (*orf02*-*orf03*) (Cheng et al., 2023; Doron et al., 2018); I55 encodes dynamins (*orf02*-*orf03*) and RecBCD (*orf23*-*orf25*); I44 encodes a classic type II restriction-modification system (*orf02*-*orf03*) (Tock & Dryden, 2005) and a DISARM system (*orf05*-*orf10*) (Ofir et al., 2017). In addition, I44 encodes a ∼14 kb operon (*orf11*-*orf20*) composed of metabolic genes including putative hydantoinases (Ishikawa et al., 1997), a peptidase M24, and an Asp/Glu/hydantoin racemase.

Given that the three elements were acquired during the course of serial transfer experiments, they must be capable of either autonomous, or non-autonomous transfer. A search of encoded genes showed no evidence of those associated with lateral transmission, such as bacterial insertion sequences (Xie & Tang, 2017), typical virus or plasmid structures (Camargo et al., 2023; de Sousa et al., 2023), or machinery for conjugation (Néron et al., 2023; M. Wang et al., 2024).

### The three GIs represent a widespread and diverse class of element in *Pseudomonas*

Given knowledge of shared features of the three elements, we turned to whole genome sequences and asked whether *Pseudomonas* genomes harbor elements with these basic features. Search criteria included the presence of *intY* downstream of *tmRNA*, as well as the presence of an *alpA*, or a 23 bp sequence (TTC AGA TTA CCC CGG CTC CAC CA, the last 14 bp being the direct repeat) that is common to the 3’-end of each element. Searches revealed 99 elements from 17,078 *Pseudomonas* genomes. Fig. 2a shows the bioinformatic workflow and Fig. 2b depicts the relationship among the elements, constructed based on previously established methods (Low et al., 2019; Nayfach et al., 2021). Briefly, 36 ORFs that are commonly present in the entire set of MGEs were selected as core genes. For each element, the core genes were concatenated in order, with gaps allowed where core genes are absent. The tree was then inferred based on the concatenated sequences. The cladogram should be interpreted in a topological sense; the edges represent evolutionary relationships only (and not distances).

**Fig. 2.**
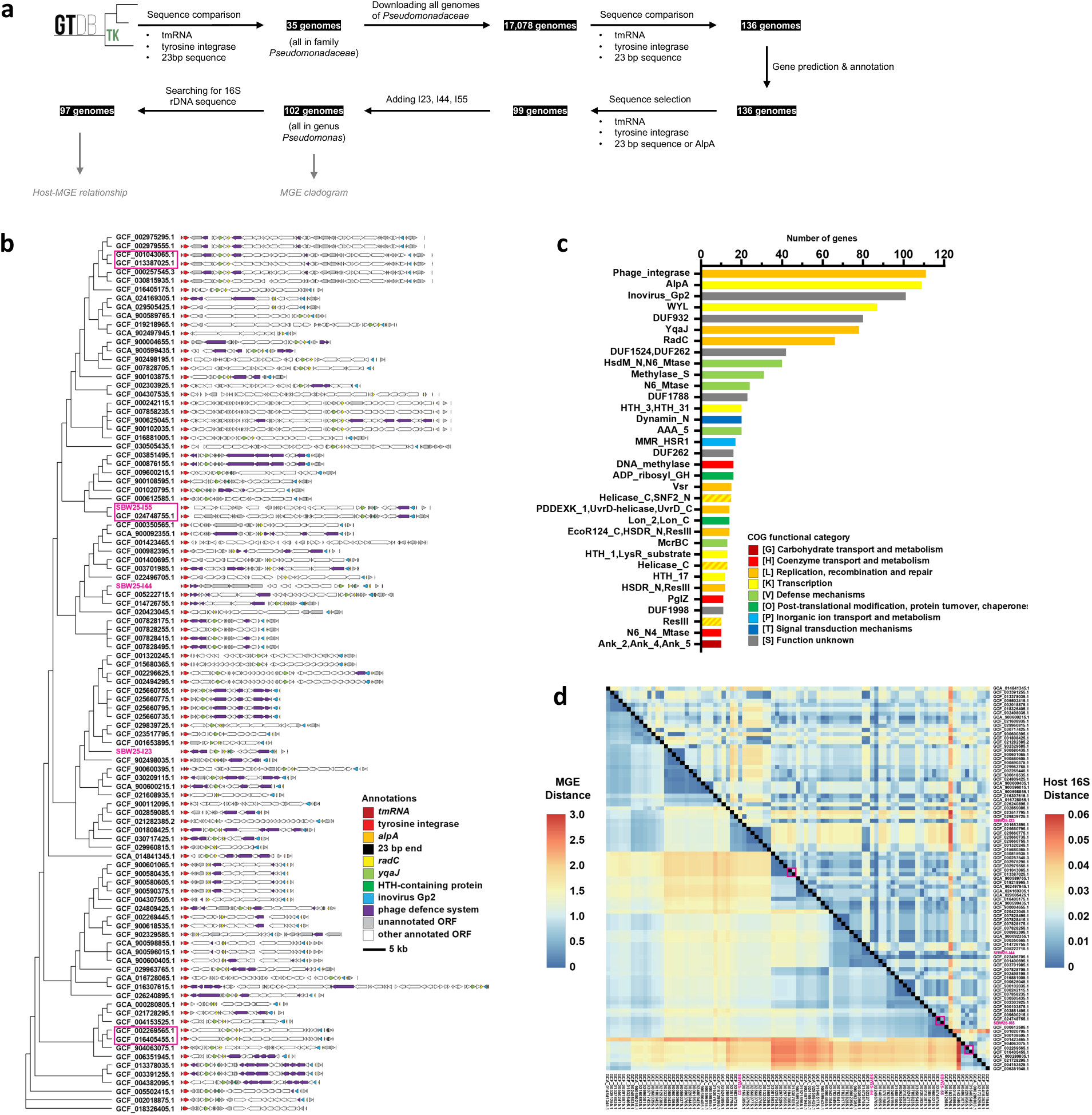
Bioinformatic analysis of similar elements. **a.** Bioinformatic pipeline for detection of similar elements to I55. Bullet points below “sequence comparison” and “sequence selection” represent the criteria for comparison and searching, respectively. **b.** Cladogram and genomic representations of I23, I44, I55 and similar MGEs (identified by NCBI RefSeq assembly indices). Basis for construction of the tree is summarized in the text. Shape of the tree represents inferred evolutionary relationship but not evolutionary distances. **c.** Rank abundance of annotated genes of different functional classes in panel b. Genes occurring at least 10 times are illustrated. A full rank abundance table is in Table S6. Colors represent COG functional categories. **d.** Pairwise distances between the MGEs (shown in lower-left), and pairwise distances between their corresponding hosts (shown in upper-right). MGEs and hosts are identified by NCBI RefSeq assembly indices. Basis for construction and interpretation of the heatmap is summarized in the text. Pink boxes highlight pairs of strains with almost identical MGE, but remarkably different 16S sequence. Annotations of those MGEs are boxed in pink in panel b.

Overall, the elements are highly diverse in terms of size and gene content (Fig. 2b). The largest elements are in excess of 70 kb with the smallest being less than 20 kb. No two elements are identical; however, several are highly similar. For example, the element detected in *P. rhodesiae* B21-046 (NCBI RefSeq Assembly GCF_024748755.1) is 99.96% identical at the nucleotide level to I55 acquired by SBW25. As described more fully below, these two host genomes are highly divergent. Other examples of such pairs are denoted by pink boxes in Fig. 2b.

Fig. 2c lists the rank abundance of genes that occur at least 10 times across the elements, along with functional classes and corresponding COG categories (Tatusov, 2000). Many elements encode homologues of RadC, YqaJ, inovirus Gp2-like protein and HTH-containing protein (Fig. 2b, 2c). These are also present in I23, I44 and I55 but were not included in the search criteria. Together, these genes, in addition to *intY* and *alpA*, likely define core element functions. In terms of functional categories, those encoding functions related to DNA replication, recombination, repair and transcription are frequently represented. Furthermore, most elements contain recognizable defense systems against bacteriophages (Fig. 2b). Many genes are found as singletons with 104 of the 241 functional gene classes identified via matches to protein databases being represented just once (Table S6). Additionally, most elements harbor at least one recognizable ORF with no matches to genes in databases (Fig. 2b).

To explore the relationship among elements and host genomes, a comparison of pairwise distances between MGEs and hosts was performed. This involved construction of phylogenetic trees for both MGEs and hosts (the latter based on 16S rDNA sequences) with the results represented in Fig. 2d. The elements shown on the X- and Y-axes are ordered based on clustering shown in Fig. 2b. The region above the diagonal represents similarity among elements, with the region below showing similarity among hosts. For example, with focus on comparison between I55 (in *P. fluorescens* SBW25) and the element in assembly GCF_024748755.1 (in *P. rhodesiae* B21-046), which is marked by a pink box: the square below the diagonal is colored dark blue indicating that the two elements are highly similar. When the same comparison is made above the diagonal, the corresponding box is light blue indicating that the host genomes carry divergent 16S rDNA sequences. Similar patterns are evident throughout; dark blue regions below the diagonal very often correspond to divergent hosts. These incongruent patterns are indicative of the transfer of similar elements across a diverse range of host genomes as expected for elements capable of frequent horizontal movement (Colombi et al., 2024; Koutsovoulos et al., 2022).

### I55 is transduced by a jumbo bacteriophage present in community filtrate

With focus on I55, we asked whether the element is capable of autonomous transfer from one SBW25 cell to another. To this end, I55 was tagged with a kanamycin resistance (Kan^R^) marker (strain MPB39465/MPB39467). The gene encoding resistance (including promoter and a synthetic transcription terminator) was inserted in the intergenic region between *orf08* and *orf09*. The location – between two convergently transcribed genes – was chosen in order to minimize chances that insertion disrupted element function. SBW25 carrying a chromosomally-integrated tetracycline resistance (Tet^R^) gene (strain MPB39500) was used as the recipient. Donor and recipient strains were co-cultured in liquid medium without shaking (Fig. 3a) and on solid agar plates (as per a standard conjugation assay). No colonies expressing both resistance markers were detected. This suggests that I55 is incapable of autonomous intercellular movement.

**Fig. 3.**
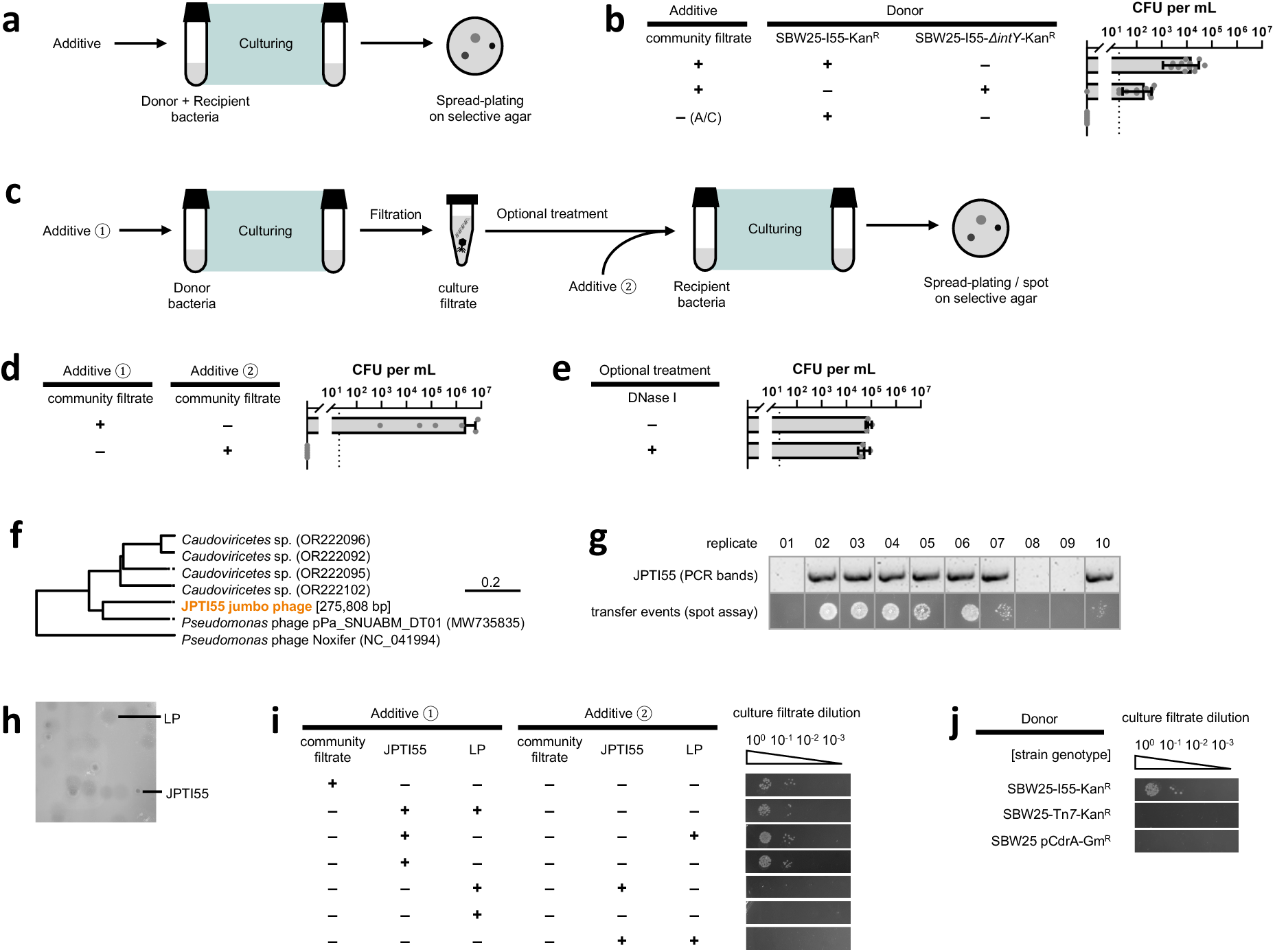
Intercellular transfer of I55. **a.** Schematic of the one-step intercellular transfer assay. **b.** The densities of CFUs on selective agar in the one-step intercellular transfer assay, to test whether and in which environment the I55 can transfer intercellularly. For all treatments, the recipient was SBW25-Tet^R^, and the final culture was spread plated on agar with Kan + Tet. Data are mean ± s.d. The dotted line represents the detection limit of the assay. The label A/C represents autoclaved filtrate. For each treatment, 8 to 12 biological replicates were used. Mann-Whitney test was applied to test the effect of IntA (*p* < 0.0001) when the community filtrate was added. **c.** Schematic of the two-step intercellular transfer assay. **d.** The densities of CFUs on selective agar in the two-step intercellular transfer assay, to test whether I55 transfers intercellular via cell-to-cell conjugation. For all treatments, the donor was SBW25-I55-Kan^R^, the recipient was SBW25-Tet^R^, no optional treatment of culture filtrate was conducted, and the final culture was spread plated on agar with Kan + Tet. Data are mean ± s.d. The dotted line represents the detection limit of the assay. For each treatment, 8 biological replicates were used. **e.** The densities of CFUs on selective agar in the two-step intercellular transfer assay, to test whether I55 transfers intercellular via transformation of naked DNA. The DNase I was confirmed to be active in digesting circular and linear DNA in the environment of community filtrate (see Fig. E3). For all treatments, the donor was SBW25-I55-Kan^R^, the recipient was SBW25-Tet^R^, additive ① was community filtrate, additive ② was none, and the final culture was spread plated on agar with Kan + Tet. Data are mean ± s.d. The dotted line represents the detection limit of the assay. For each treatment, 3 biological replicates were used. Mann-Whitney test was applied to test the effect of DNase I (*p* = 0.4000) in transfer. **f.** A phylogeny of relatives of the jumbo bacteriophage JPTI55. The complete genome of JPTI55 was *de novo* assembled from metagenomic sequencing, later corrected by whole genome sequencing of phage isolates. The genome of JPTI55 is circular and 275,808 bp in length. Distance on the phylogeny represents identity in nucleotide sequences. **g.** The relationship between the presence of JPTI55 and the presence of transfer events. Two-step intercellular transfer assay was conducted. For all treatments, the donor was SBW25-I55-Kan^R^, the recipient was SBW25-Tet^R^, additive ① was a certain dilution level of community filtrate, additive ② was none, no optional treatment of culture filtrate was conducted, and the final cultures were independently spotted on agar with Kan + Tet. There were 10 technical replicates for each type of additive ①. In addition, the final cultures were filtrated and tested for the presence of JPTI55 by PCR. Picture here corresponds to the set of 10 replicates receiving a certain dilution level of community filtrate as additive ①, at which around half of the replicates contained JPTI55 and the other half did not. Photos of five additional sets of replicates, receiving community filtrate from another dilution level or from another biological replicate of community filtrate, can be found in Fig. S1. **h.** Typical photos of the plaque assay of the community filtrate against SBW25 wild type as hosts, using 0.2% top agar in M9+Glucose medium. Picture contrast was altered to facilitate visualization of the plaques. The identities of the plaques were confirmed by PCR. **i.** The presence or absence of transfer events along a dilution series of culture filtrate in the two-step intercellular transfer assay, to test the transfer ability of the UFV-like bacteriophage and the JPTI55 jumbo bacteriophage. For all treatments, the donor was SBW25-I55-Kan^R^, the recipient was SBW25-Tet^R^, at the optional treatment step the culture filtrate was diluted along a dilution series and each dilution was separately added to independent recipient cultures, and the final cultures were independently spotted on agar with Kan + Tet. Picture contrast was altered to facilitate visualization of the spots. Photos of two additional biological replicates can be found in Fig. S2. **j.** The presence or absence of transfer events along a dilution series of culture filtrate in the two-step intercellular transfer assay, to test whether JPTI55 could mobilize chromosomal markers or plasmids. For all treatments, the recipient was SBW25-Tet^R^, additive ① was JPTI55, additive ② was none, at the optional treatment step the culture filtrate was diluted along a dilution series and each dilution was separately added to independent recipient cultures, and the final cultures were independently spotted on agar with Kan + Tet (except for the treatment with donor strain SBW25 pCdrA-Gm^R^, which was spotted on agar with Gm + Tet). Picture contrast was altered to facilitate visualization of the spots. Photos of two additional biological replicates can be found in Fig. S3.

In the original serial transfer experiment, filtrate from serially-propagated compost C had been added to SBW25 cultures. We thus considered it possible that one or more components of the filtrate might be necessary to mobilize the element. To this end, the experiment above was repeated in liquid medium, but this time with the addition of filtrate from compost C (Fig. 3a). On plating mixtures to selective agar an average ∼10^4^ colonies (per mL) resistant to kanamycin and tetracycline were detected (Fig. 3b). Deletion of the tyrosine integrase *intY* encoded on I55, resulting in donor strain SBW25-I55Δ*intY*-Kan^R^ (strain MPB39565/MPB39568), significantly reduced the double resistant colonies to ∼10^2^ colonies (per mL) (*p* < 0.0001). This indicates that circularization – or linear concatenated repeats – of I55 (see Fig. E2 for details) underpins intercellular transfer.

Next, we asked whether donor-recipient cell-cell contact is necessary for the transfer of I55. To test this, the transfer process was divided into two steps (Fig. 3c). In the first step, SBW25-I55-Kan^R^ with filtrate from compost C was cultured and the filtrate collected. In the second step, recipient SBW25-Tet^R^ was grown with addition of the filtrate obtained from the first step. The mixture was plated as previously. On average, more than 10^6^ colonies (per mL) resistant to both antibiotics were detected (Fig. 3d), indicating that transfer of I55 is independent of cell-cell contact. However, if the compost filtrate was added at the second step and not the first, then no double-resistant colonies were detected (Fig. 3d). I55 therefore requires some component in the community filtrate in order to mediate transfer to recipient cells.

While phages are plausible facilitators of horizontal transfer, there existed the possibility that phages may be indirectly involved and required solely for cell lysis. If so, then I55 would exist as naked DNA during transfer and susceptible to degradation by DNase. Culture filtrate applied to the recipient culture was therefore treated with DNase I, but no reduction in the number of transformants was observed (Fig. 3e).

To identify the filtrate component necessary for mobilization, serial dilutions of the community filtrate were made with the aim of diluting out the transfer agent. A sample from each dilution was then used as additive ① in the two-step transfer assay (Fig. 3c). In each instance a threshold was observed below which no transfer occurred. Samples immediately above and below the critical limit were subject to amplification and DNA sequence analysis. The resulting DNA sequence reads revealed presence of a jumbo phage similar to *pPa_SNUABM_DT01* (Fig. 3f) and hereafter referred to as JPTI55. To determine the correlation between presence of the JPTI55 and I55 transfer, the two-step transfer assay exploiting the dilution-to-extinction design was conducted for three independent filtrate samples, each with 10 biological replicates at each dilution. Transfer of I55 was observed if and only if JPTI55 was detected (by PCR) in the filtrate (Fig. 3g).

Next, we sought to isolate JPTI55 and thus confirm by direct experimentation the involvement of JPTI55 in transfer. To this end aliquots of community filtrate from various dilutions were plated against SBW25. On standard 0.4% top-agar LB plates, morphologically uniform large plaques (LP) were detected, but these plaques were present at dilutions lower than those that resulted in I55 transfer. Moreover, PCR analysis of plaques using primers diagnostic for JPTI55 proved negative. Attempts to transfer I55 to new hosts via addition of phages purified from these plaques also proved negative.

The experiment was therefore repeated but plating was performed on SBW25 contained in 0.2% top-agar M9-glucose plates. At this reduced agar concentration – and at dilutions corresponding to those in which JPTI55 was known to be present in the filtrate – two morphologically distinct plaques were observed. PCR analysis confirmed that the small clear (less abundant) plaques (see Fig. 3h) contained JPTI55.

Individual plaques were subject to three rounds of purification by plaque assay. Finally, the two-step transfer assay was conducted with JPTI55 and/or LP phages added to donor or recipient cultures. The results (Fig. 3i) demonstrate that JPTI55 was solely responsible for the transfer of I55 – and did so as efficiently as the culture filtrate from which it was isolated.

To directly test whether I55 behaves as a phage satellite, we assessed whether the jumbo phage JPTI55 could non-specifically package host DNA. If I55 were transferred simply as part of generalized transduction, chromosomal markers unrelated to I55 would also be detected among transductants. We engineered SBW25 strains carrying chromosomal (strain MPB29284) and plasmid-borne (strain MPB16791) antibiotic resistance markers, and repeated the two-step transfer assays with purified JPTI55. No transfer of chromosomal or plasmid markers was observed (Fig. 3j), indicating that generalized transduction does not occur (or is below the detection limit). These findings suggest that I55 is preferentially packaged and transferred by JPTI55 in a manner consistent with the behavior of a phage satellite (Penadés et al., 2025).

### A type II restriction-modification system protects against UFV-like phages

We next sought understanding of the fitness consequences of element carriage and again focused attention on I55. I55 encodes several putative phage defense systems (*orf02*, *orf17*, *orf24*, *orf37*), and compost filtrate is rich in phages (Quistad et al., 2020). We therefore hypothesized that I55 confers a fitness benefit to its host by providing resistance against phages. By fluorescence-based competition assays (Fig. 4a), the data show that I55 confers a significant fitness advantage (∼6%) in the presence of compost filtrate (*p* = 0.0104), but is costly (∼2%) in the absence of the filtrate (*p* = 0.0033) (Fig. 4b). To directly test the hypothesis that the benefit of I55 carriage arises from phage protection, compost C filtrate was plated against lawns of SBW25 and of SBW25 carrying I55. Plaques were observed on both hosts, but carriage of I55 significantly reduced infection and cell lysis (Fig. 4c).

**Fig. 4.**
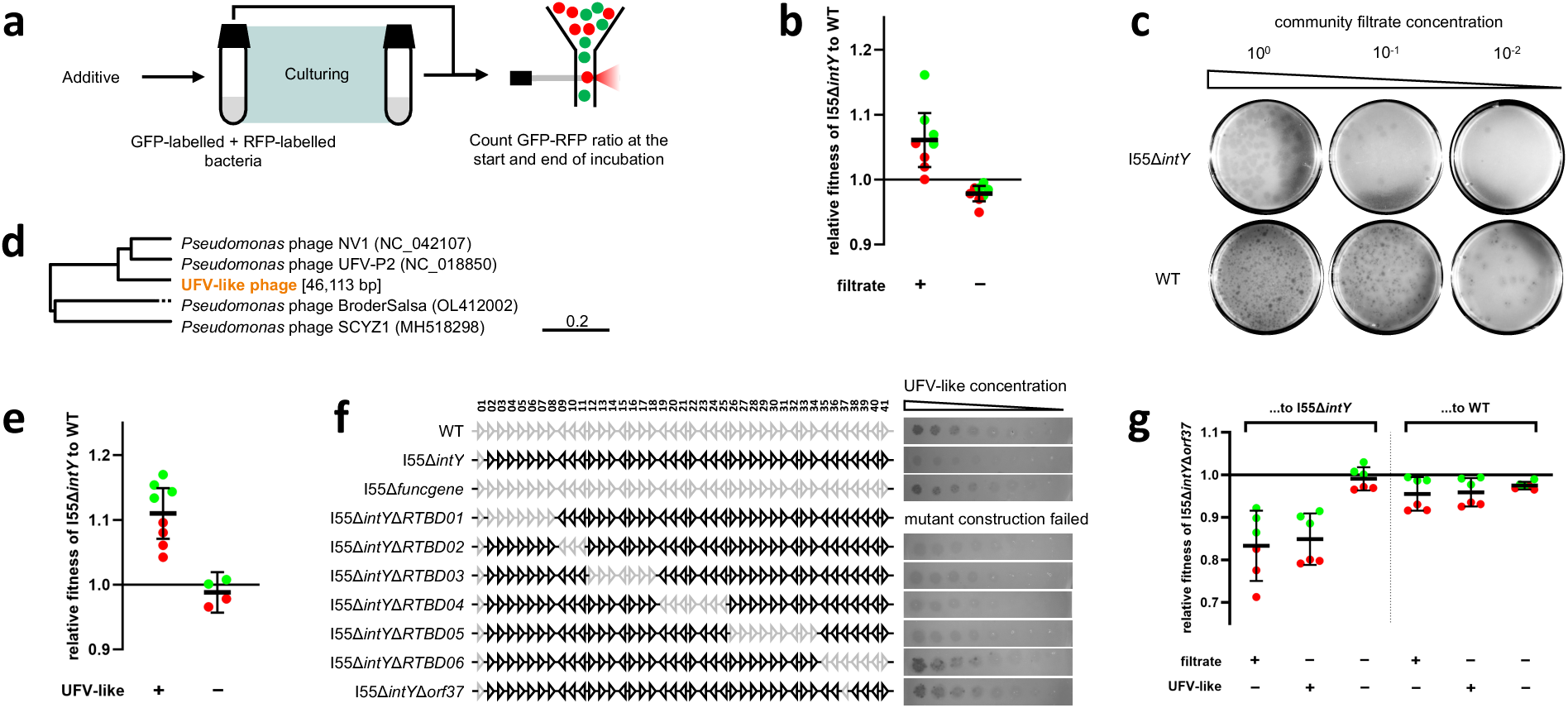
Fitness advantage and phage defense activity of I55. **a.** Schematic of fluorescent fitness assay. To eliminate the possibility for I55 to transfer among cells during the fitness assay, all strains carrying I55 had *intY* deleted, because SBW25-I55Δ*intY* was previously shown to be incapable of excision from the genome (see Fig. E2). Moreover, to ensure the two competing strains have identical genetic backgrounds, ancestral SBW25 was re-constructed by deleting I55 from SBW25-I55Δ*intY*. **b.** Fluorescent fitness assay results for SBW25-I55Δ*intY* (strain MPB39132/MPB39136) relative to wild type (strain MPB39379/MPB39459), in presence or absence of community filtrate. Data are mean ± 95% CI. In the Filtrate + and - treatments, stored filtrate aliquots from compost C and from SBW25 were added, respectively. The color of each data point corresponds to the color of the fluorescent marker added to SBW25-I55Δ*intY*. For each treatment, 8 biological replicates (4 for either tagging strategy) were used. One sample *t*-tests were applied to test the relative fitness values against 1 when the community filtrate was added (*p* = 0.0104) and not added (*p* = 0.0033). **c.** Typical photos of the plaque assay of the community filtrate against SBW25-I55Δ*intY* or wild type as hosts. Picture contrast was altered to facilitate visualization of the plaques. Photos of three additional biological replicates can be found in Fig. S4. **d.** A phylogeny of relatives of the UFV-like bacteriophage. The complete genome of the UFV-like phage was *de novo* assembled from whole genome sequences. The genome of UFV-like phage is circular and 46,113 bp in length. Distance on the phylogeny represents identity in nucleotide sequences. **e.** Fluorescent fitness assay results for SBW25-I55Δ*intY* relative to wild type, in presence or absence of the UFV-like bacteriophage. Data are mean ± 95% CI. The color of each data point corresponds to the color of the fluorescent marker added to SBW25-I55ΔintY. For each treatment, 8 or 4 biological replicates (4 or 2 for either tagging strategy, respectively) were used. One sample *t*-tests were applied to test the relative fitness values against 1 when the UFV-like bacteriophage was added (*p* = 0.0003) and not added (*p* = 0.3118). **f.** Systematic deletion assay and phage spot assay to identify the phage defense gene. Regions of I55 deleted are shown in grey. Picture contrast was altered to facilitate visualization of the spots. Photos of two additional biological replicates can be found in Fig. S5. **g.** Fluorescent fitness assay results for SBW25-I55Δ*intY*Δ*orf37* (strain MPB42855) relative to SBW25-I55Δ*intY* or wild type, in presence or absence of the community filtrate or UFV-like bacteriophage. Data are mean ± 95% CI. The color of each data point corresponds to the color of the fluorescent marker added to for SBW25-I55Δ*intY*Δ*orf37*. For each treatment, 6 biological replicates (3 for either tagging strategy, respectively) were used. One sample *t*-tests were applied to test the relative fitness values of SBW25-I55Δ*intY*Δ*orf37* relative to SBW25-I55Δ*intY* against 1 when the community filtrate was added (*p* = 0.0035), when the UFV-like bacteriophage was added (*p* = 0.0014), and when there was no additive (*p* = 0.4237). The same tests were applied to test the relative fitness values of SBW25-I55Δ*intY*Δ*orf37* relative to wild type against 1 when the community filtrate was added (*p* = 0.0331), when the UFV-like bacteriophage was added (*p* = 0.0252), and when there was no additive (*p* = 0.0008).

Five phages – all with the LP morphology – were purified from single plaques on plates shown in Fig. 4c. Whole genome sequencing revealed the phages to be identical at the nucleotide level. Interrogation of the phage genome against NCBI databases showed that the closest relative is *Pseudomonas* phage UFV-P2 (Fig. 4d). Henceforth we refer to phages from large plaques as UFV-like phages. The fluorescence-based competition assay was then repeated with pure UFV-like phages, and the assay result was qualitatively reproduced (see Fig. 4e). This shows that defense against UFV-like phage present in the community filtrate partially explains the fitness benefit of I55.

To determine regions of I55 responsible for protection against the UFV-like phages, the element I55 was divided into six regions (RTBD01-06, see Fig. 4f) according to operon structures, and each region was separately deleted (strains MPB42170, MPB41585, MPB42372, MPB42031 and MPB41438, for Δ*RTBD02*, Δ*RTBD03*, Δ*RTBD04*, Δ*RTBD05* and Δ*RTBD06*, respectively). Spot assay with UFV-like phages revealed that deletion of the sixth region (RTBD06) abolished the phage defense phenotype of I55, while deletions of the other five regions (except RTBD01, which we were unable to construct) maintained defense level of intact I55 (Fig. 4f). DefenseFinder predicts *orf37*, encoded within region RTBD06, to be a type II-G restriction-modification system. We thus deleted *orf37* – producing SBW25-I55Δ*intY*Δ*orf37* (strain MPB42855) – and repeated the spot assay, and observed infection levels comparable to SBW25. This confirmed that the phage defense phenotype of I55 is encoded by *orf37*.

Finally, we tested whether *orf37* is indeed the cause of the fitness benefit of I55 in environments with community filtrate or UFV-like phages. Fluorescence-based competition assays were conducted with SBW25-I55Δ*intY*Δ*orf37* against SBW25-I55Δ*intY*. As expected, deletion of *orf37* significantly reduced host fitness in presence of community filtrate or UFV-like phages (*p* = 0.0035 and 0.0014, respectively, see Fig. 4g), but had no fitness influence in the absence of both (*p* = 0.4237), which supported the hypothesis that the fitness effect of I55 is explained by *orf37*. Competition of SBW25-I55Δ*intY*Δ*orf37* against the wild type SBW25 showed that the former was always slightly less fit than the latter regardless of the environment (*p* = 0.0331, 0.0252 and 0.0008, in environments with community filtrate, with UFV-like phages and with no additives, respectively).

## Discussion

Horizontal gene transfer (HGT) mediated by mobile genetic elements (MGEs) shapes the diversity and function of microbial communities, but much remains to be discovered regarding both the vehicles of transfer and the functional genes subject to horizontal movement. The study reported here, was motivated by earlier work (Quistad et al., 2020), in which repeated exposure of compost community filtrates to fresh bacterial hosts caused a substantial fraction of community DNA to enter the horizontally disseminated pool. Comparative metagenomics revealed genes of ecological significance and, in some cases, known MGEs such as phages and plasmids. However, progress has been constrained by the inability to experimentally capture unknown elements in culturable hosts.

Here, we simplified the original strategy: we retained the complexity of community-derived filtrates but exposed these to a single, genetically tractable strain of *Pseudomonas fluorescens*. Within a short period, the focal bacterium acquired three MGEs – designated I23, I44, and I55 – each integrating immediately downstream of the *tmRNA* gene, but lacking typical structures indicative of autonomous movement (Camargo et al., 2023; de Sousa et al., 2023; Néron et al., 2023; M. Wang et al., 2024; Xie & Tang, 2017). Using minimal defining features – target site, 3ʹ-end sequence, and integrase – we identified ∼100 related elements within *Pseudomonas*, though the class is broader. Relaxing constraints on flanking sequence similarity revealed that elements with essential features of the three MGEs are widespread. Many such genomic islands insert at *tmRNA* or *tRNA* genes, are flanked by direct repeats, encode tyrosine recombinases, and lack genes for autonomous transfer. The “phage defense elements” recently described by (Hussain et al., 2021) exemplify this class. Moreover, Islander (Hudson et al., 2015), a tool for detecting integrases downstream of *tmRNA*/*tRNA*, shows that similar elements are present across both eubacteria and archaea.

Starting from evidence that I55 is incapable of autonomous transfer, we hypothesized the involvement of a helper element (Ares-Arroyo et al., 2024). The occurrence of I55 transfer in the absence of cell–cell contact and despite DNase treatment pointed to a phage-mediated mechanism. Using dilution-to-extinction assays on filtrates, we linked transfer to the presence of a jumbo bacteriophage, JPTI55. Further experiments revealed that I55 is transduced by JPTI55 as a satellite – a finding that, to our knowledge, is the first report of a phage satellite parasitizing a jumbo phage.

The SBW25–I55–JPTI55 system opens new avenues for exploring unconventional mechanisms of gene transfer and for developing tools for genetic engineering. Most known phage satellites parasitize phages with genomes under 200 kb (Penadés et al., 2025), whose replication and packaging occur in the cytoplasm. In contrast, jumbo phages form proteinaceous shells that enclose replicating DNA in the cytoplasm (Laughlin et al., 2022), potentially creating a barrier to satellite hijacking. That I55 can bypass or penetrate this structure suggests novel biology. Additionally, the known size constraints of standard phage capsids limit satellites to ∼20 kb, restricting their use as vectors. With I55’s larger genome and its association with the large JPTI55 capsid, the possibility arises for shuttling larger cargo between cells. However, identifying the minimal satellite backbone, host range and methods for phage elimination is a prerequisite for realizing this potential.

Beyond I55’s mode of transfer, its experimental capture provides a paradigm for uncovering MGE mobility in complex environments. A similar approach may help resolve the mystery surrounding the ∼41 kb *Vibrio* Pathogenicity Island-1 (VPI-1), which encodes the toxin-coregulated pilus receptor for CTXΦ, the cholera toxin phage. VPI-1 acquisition enables subsequent CTXΦ entry and conversion of environmental *Vibrio cholerae* into toxigenic strains (Faruque & Mekalanos, 2003, 2012). Although VPI-1 is known to have transferred horizontally (Karaolis et al., 1999; O’Shea & Boyd, 2002), its mobility remains controversial (Faruque et al., 2003). A protocol akin to that used to identify JPTI55 may lead to identification of a secondary transfer agent – possibly also a jumbo phage – providing a strategy to block transmission.

A benefit of working with I55 in a genetically tractable host was the ability to determine fitness effects. Acquisition of I55 conferred a clear advantage in the presence of compost filtrate (but a small cost in its absence), explaining its detection in the limited number of sequenced colonies arising from the initial selection experiment. The benefit appears to stem in part from phage defense. Experiments with UFV-like phages purified from filtrate confirmed I55’s protective effect. Systematic deletions identified *orf37*, a type II restriction-modification system, as responsible for resistance to UFV-like phages. Other predicted phage defense systems in I55 – including dynamin and RecBCD orthologs (Dillingham & Kowalczykowski, 2008; Guo et al., 2022) – were not active against these phages, but could function in other contexts. More generally, this finding supports the view that MGEs are reservoirs of phage defense systems, which enhance their own propagation (Bernheim & Sorek, 2020; Hussain et al., 2021; Johnson et al., 2023; Rousset et al., 2022).

The discovery of I55-family elements and their JPTI55-mediated mobilization exposes a previously hidden layer of HGT involving jumbo phages and satellites. While the elements described here represent only a fraction of the diversity likely present, they highlight how microbial communities can facilitate gene flow at scales reminiscent of sexual reproduction in eukaryotes (Rainey & Quistad, 2020). Applying this experimental strategy to other hosts and habitats is likely to uncover further mobile elements, novel gene cargos, and distinct transfer mechanisms – critical for understanding the ecological and evolutionary forces that shape microbial communities.

## Materials and Methods

### Strains and growth conditions

All bacterial strains used in this study are listed in Table S1. All strains were preserved in -80 °C with 30% glycerol saline.

*Isolation of foreign elements, plasmid construction, strain construction*: Unless otherwise stated, *P. fluorescens* SBW25 strains were cultured in LB-Miller (LB) broth at 28 °C, while *E. coli* strains were cultured in LB (with antibiotics where maintenance of plasmids was required) at 37 °C, both shaking at 220 rpm for 16-24 h.

*Experimental evolution, intercellular transfer assay and fluorescent fitness assay*: Unless otherwise stated, each pre-conditioning and incubation was performed in the absence of shaking at 28 °C, in 5 mL freshly made M9-glucose medium (M9 salt solution supplemented with 2 mM MgSO4, 0.1 mM CaCl2 and 0.2% w/v D-glucose) in a 15 mL Falcon tube.

*Plaque assay and bacteriophage isolation*: All assays were done in S-LB environment, except the assay for isolation of the jumbo phage JPTI55 (see below) which was done in M9-glucose environment. For the S-LB environment, unless otherwise stated, bacterial host strains were cultured in S-LB medium (LB medium supplemented with 10 mM MgCl_2_ and 5 mM CaCl_2_) at 28 °C, shaking at 220 rpm for 16-24 h until stationary phase. The semi-soft agar (SSA) used consisted of S-LB medium and 0.4% agar. The bottom agar consisted of LB medium and 1.5% agar. For the M9-glucose environment, unless otherwise stated, bacterial host strains were cultured in M9-glucose medium at 28 °C, shaking at 220 rpm for 16-24 h until stationary phase. The semi-soft agar (SSA) used consisted of M9-glucose medium and 0.2% agar. The bottom agar consisted of M9-glucose medium and 1.5% agar.

Bacterial culture was plated on respective growth medium supplemented with 1.5% agar. Unless otherwise stated, agar plates were incubated at 28 °C. Antibiotics were applied as per the following concentrations: kanamycin (Kan, 50 μg/mL), nitrofurantoin (100 μg/mL), tetracycline (Tet, 12.5 μg/mL), gentamicin (Gm, 20 μg/mL), *Pseudomonas* CFC selective supplement (Millipore, as per the manual).

### Compost community sampling, storage and retrieval

Three heaps of garden compost, named B, C and D, were sampled within 10 km of Plön, Germany (54°9ʹ44ʺN, 10°25ʹ17ʺE), from May to August 2022. Upon collection, the samples were immediately transferred to laboratory and processed as following. For each sample, 30 g compost was mixed with 150 mL M9 salt solution, shaken vigorously for ∼30 sec, and left static for ∼5 min. The supernatant was distributed into 1 mL aliquots, to each of which 350 μL glycerol saline was added. The aliquots were frozen in -80 °C for storage.

To retrieve the compost wash, one aliquot from the frozen stock was kept on ice for ∼10 min. After thawing, the mixture was pelleted by centrifuging at 15000 rcf for 1 min, resuspended in 1 mL M9 salt solution, and then pelleted by centrifuging again. The washed pellet was then resuspended in 500 μL M9 salts solution to inoculate microcosms.

### Community filtrate (fixed) generation, storage and retrieval

Initially, 500 μL washed suspension from one compost wash aliquot was inoculated into 100 mL M9-glucose medium in a 500 mL microcosm. The microcosm was incubated without shaking at 28 °C for 7 days. At the end of incubation period, the whole culture was filtered with a 0.2 μm filter (Whatman). The filtrate was distributed into 600 μL aliquots. Aliquots were directly frozen at -80 °C for storage.

To retrieve the community filtrate, one aliquot from the frozen stock was kept at room temperature for ∼10 min. After thawing, the aliquot was directly used to inoculate microcosms. Unless otherwise stated, 50 μL community filtrate was added to each microcosm.

As a negative control, *P. fluorescens* SBW25 was used in place of the washed suspension, and otherwise followed exactly the same generation, storage and retrieval protocols to obtain the SBW25 filtrate to inoculate microcosms.

### Experimental evolution

#### Co-culture regime

Parallel lines of SBW25, cocultured with the compost community, were serially propagated. In detail, SBW25-RFP-Gm^R^ (strain MPB28806) and 90 μL washed suspension from compost C were inoculated. There were three parallel lines, with one control line where no washed suspension was added. For each line, an 1/20 transfer was made every 24 h, for a total of five transfers. At each transfer, a dilution series of the culture was spread plated on LB + CFC + Gm plates, which were then cultivated for 24 h. The number of RFP colonies were counted and recorded.

#### Fixed filtrate regime

Parallel lines of SBW25 were serially propagated, with aliquots of filtrate from the frozen stock added at each transfer. In detail, filtrate from communities B, C and D, and from SBW25, were separately used. For each type of filtrate, there were five parallel lines of SBW25-GFP-Kan^R^ (strain MPB29284) and five parallel lines of SBW25-RFP-Kan^R^ (strain MPB29300). For each line, an 1/100 transfer was made every 48 h, for a total of seven transfers. At each transfer, 50 μL filtrate was added to a fresh microcosm, and aliquots from the old microcosm were stored at -80 °C.

#### Live filtrate regime

Parallel lines of SBW25 were serial propagated, with aliquots of filtrate from independently propagated communities or SBW25 added at each transfer. In detail, 200 μL washed suspension from compost C wash was added and transferred 1/100 every 48 h, yielding fresh filtrate by filtering 1mL of the culture through a 0.2 μm filter (Whatman). As a negative control, *P. fluorescens* SBW25 was used in place of the washed suspension, but otherwise the same protocol was followed to obtain SBW25 filtrate. At the same time, for each type of filtrate, there were four parallel lines of SBW25-GFP-Kan^R^ (strain MPB29284) and four parallel lines of SBW25-RFP-Kan^R^ (strain MPB29300). For each line, a 1/100 transfer was made every 48 h, for a total of 14 transfers. At each transfer, 50 μL fresh filtrate was added to the new microcosm, and a sample from the old microcosm were stored at -80 °C.

### PCR and gel electrophoresis

All PCR primers used in this study are listed in Table S3. PCR was done with NEB Q5 polymerase, with either extracted DNA or bacterial suspension as templates. Annealing temperatures and extension time was determined as per NEB Tm Calculator (version 1.16.6). Unless otherwise stated, 35 cycles were performed. Gel electrophoresis was performed for 5 μL PCR products with 1x DNA loading dye, in 1% agarose gels.

### Detection, isolation and annotation of foreign elements

For each line, frozen culture at the last transfer was serially diluted and plated out on LB plates. For the live-filtrate regime, 2 to 3 colonies were picked and separately cultured overnight, from which aliquots were frozen and DNA extracted with Qiagen DNeasy UltraClean Microbial kit. DNA samples were sequenced by Illumina NextSeq for at least 300x coverage. For fixed-filtrate regime, 24 colonies were randomly picked and cultured up overnight in 300 μL LB in 96-deep-well plates. Aliquots from each of the 24 cultures were mixed in equal proportion and then DNA extractions were performed with Qiagen DNeasy UltraClean Microbial kit. The leftovers in 96-deep-well plates were stored at -80 °C. DNA samples were Illumina short-read sequenced (read length 150 bp) by Eurofins Scientific SE for at least 300x coverage.

Bioinformatic analysis of sequencing data was done in Geneious Prime 2022.2.1 (https://www.geneious.com). For each sample, raw reads were trimmed by BBDuk and mapped to *P. fluorescens* SBW25 reference genome (Fortmann-Grote et al., 2023) under the default settings. The unmapped reads were saved and then *de novo* assembled. The vast majority of contigs were short (< 1 kb) and apparently artefacts of sequencing technology. However, in some samples, long (> 10 kb) and good quality contigs were obtained. To test if the contig inserted into the genome as a foreign element, the first 50 bp and last 50 bp of the contig were blasted against the reference genome. If the two regions mapped, manual comparison were used to confirm the insertion and its locus.

To confirm the insertion of the foreign element, two primer pairs were designed, flanking either the left or the right junction of the insertion. For fixed-filtrate regime, the two primer pairs were additionally used to exclude the cultured isolates (out of the 24 picked) that carried the foreign element. Such isolates were then routinely whole-genome sequenced by Eurofins Scientific SE and Variations / SNPs calling was subsequently performed. Table S5 shows the list of mutations detected in each isolate.

The three foreign elements were screened for open reading frames (ORFs) by Prokka (Seemann, 2014) in KBase (Arkin et al., 2018). The ORFs were then annotated by Pfam in InterProScan (Jones et al., 2014). The elements were also screened by DefenseFinder (Tesson et al., 2022) for potential phage defense systems. Alignment of the 3 foreign elements was done by Easyfig (M. J. Sullivan et al., 2011) based on blastn under the default settings.

### Detection and analysis of similar elements in databases

The nucleic acid sequences of the *tmRNA*, *intY* (from the element I55) and the 23 bp sequence at the 3’- end were compared to representative genomes in GTDB-tk (version R214) (Chaumeil et al., 2020). Sequence comparisons revealed that 35 representative genomes contain all three sequences, and they all belong to the family *Pseudomonadaceae*. This motivated a more focused and thorough search within this family. Therefore, the nucleic acid sequences of the *tmRNA*, *intY* (from the element I55) and the 23 bp sequence at the 3’-end were compared to all available genomic sequences (17,078 in total) of strains belonging to the family *Pseudomonadaceae*. Sequence comparisons revealed that 136 genomes in the family *Pseudomonadaceae* contain all three sequences. Those 136 genomes were then subject to gene prediction by Prodigal (v2.6.3) (Hyatt et al., 2010) and functional annotation by eggNOG-mapper (v2) (Cantalapiedra et al., 2021). Based on annotations, 99 genomes fulfilled the following criteria: (1) contained an annotated tmRNA; (2) contained an annotated tyrosine integrase (IntY); (3) contained at least one of an annotated AlpA or the 23 bp sequence. These 99 genomes are derived from 67 species, all in the genus *Pseudomonas*. In each genome, the MGEs were isolated by extracting the sequence from the 3’-end of the *tmRNA* to the 3’-end of the 23 bp sequence or the 3’-end of *alpA*. All MGEs were screened by DefenseFinder (Tesson et al., 2022) for potential phage defense systems.

To establish evolutionary relationships of the 99 detected MGEs and I23, I44 and I55, a cladogram of the 102 MGEs was built based on previously established methods (Low et al., 2019; Nayfach et al., 2021). Briefly, 36 ORFs that are commonly present in the MGEs were selected as marker genes. For each MGE, the marker genes were concatenated in order, with a gap introduced in cases where a marker gene was absent. The tree of the MGEs was then inferred based on the concatenated sequences.

To assess host-MGE relationships, the 16S rDNA sequences of the hosts of the 102 MGEs were extracted. Among them, 5 host genomes lack detectable 16S rDNA regions and hence were excluded from the following analysis. For the 97 remaining genomes, phylogenetic trees were built separately based on the 16S rDNA sequences by Clustal (v2.1) (Sievers et al., 2011), and based on the MGE sequences by the same method as before (Low et al., 2019; Nayfach et al., 2021). The two trees were converted to distance matrices in R (version 4.1.2), with rows clustered according to MGE distances.

### Plasmid construction

All plasmids used in this study are listed in Table S2. Plasmid pUIsacB (plasmid MPB35522) backbone was amplified by PCR. SBW25 genomic fragments flanking the region of interest, where a mutagenesis, knock-out or knock-in was to be performed, were amplified by PCR. In particular, the Kan^R^ fragment used to tag I55 was produced by cloning the Kan^R^ gene from plasmid pMRE-Tn*7*-152 (plasmid MPB27708) followed by a synthetic transcription terminator (CCA ATT ATT GAA CAC CCT TCG GGG TGT TTT TTT GTT TCT GGT CTC CC). Primers were designed with a 5’ overhang to facilitate assembly. Backbone and fragments were assembled (with NEBuilder HiFi DNA Assembly Master Mix) and transformed into *E. coli* Top10 competent cells. Successful transformants were selected on LB + Tet plates, and confirmed by PCR and Sanger sequencing across the construct.

### Strain construction

#### Two-step allelic replacement (2SAR)

The method is adapted from Hmelo et al. (Hmelo et al., 2015) and extensively described by Farr et al. (Farr et al., 2025). Briefly, recipient (*P. fluorescens* SBW25), donor (*E. coli* carrying the plasmid used for allelic exchange) and helper (*E. coli* carrying plasmid pRK2013, plasmid MPB35623) strains were cultured overnight until stationary phase. Then, 1 mL recipient culture was heat-shocked at 42 °C for 20 min, and subsequently mixed with 500 μL donor and 500 μL helper culture. The mixture was pelleted and resuspended in 100 μL LB, which was then spotted onto an LB plate for incubation overnight. The cultivated cell layer was then plated out on an LB + nitrofurantoin + Tet plate and incubated overnight to select for trans-conjugates. Single trans-conjugates were then picked and streaked onto TYS10 plates, which were subsequently incubated overnight. Finally, PCR and Sanger sequencing were used to screen for the desired mutants.

#### Fluorescent tagging

The method is adapted from Schlechter et al. (Schlechter et al., 2018). Briefly, recipient (*P. fluorescens* SBW25) and donor (*E. coli* carrying the plasmid pMRE-Tn*7*-152 or pMRE-Tn*7*-155, MPB27708 or MPB27709, respectively) strains were cultured overnight until stationary phase. Then, 1 mL recipient culture was heat-shocked at 42 °C for 20 min, and subsequently mixed with 1 mL donor culture. The mixture was pelleted and resuspended in 100 μL LB, which was then spotted onto an LB plate for incubation overnight. The cultivated cell layer was then plated out on an LB + nitrofurantoin + Kan + 0.1% w/v arabinose plate and incubated overnight to select for trans-conjugates. Single trans-conjugates were picked, and integration of the fluorescent tag was then confirmed by PCR.

#### *Preparation of* SBW25-I55 *and* SBW25 *(reconstructed) for assays*

DNA sequencing of the evolved isolate carrying I55 (strain MPB33357) revealed, in addition to presence of the mobile element, two SNPs elsewhere in the genome (see Table S5) that had arisen during the course of serial propagation. Additionally, this isolate also harbored the Tn*7*-152 cassette. While there was no reason to suspect any connection to I55, these SNPs were reverted to the ancestral SBW25 sequence, and Tn*7*-152 removed, by site-directed mutagenesis using the 2SAR strategy described above. Prior to these mutagenesis steps, two independent colonies were picked from the evolved isolate and treated as biological replicates in all subsequent genetic manipulations. The resulting genotype, SBW25-I55 (strain MPB39024/MPB39026) is in principle isogenic with SBW25 (strain MPB29212). However, to be more certain of this fact, the ancestral SBW25 genotype used in the fitness assays and plaque assays was re-generated from SBW25-I55 by removal of the element, giving reconstructed SBW25 (strain MPB39379/MPB39459).

### Intercellular transfer assay

#### Conjugation regime

The donor, a SBW25 strain with Kan^R^-tagged I55 (strain MPB39465/MPB39467), recipient, a SBW25 strain with a Tet^R^ marker (strain MPB39500), and helper, *E. coli* carrying plasmid pRK2013 (plasmid MPB35623), were cultured overnight until stationary phase. Then, 1 mL recipient culture was heat-shocked at 42 °C for 20 min, and subsequently mixed with 500 μL donor and 500 μL helper culture. The mixture was pelleted and resuspended in 100 μL LB, which was then spotted onto an LB plate for incubation overnight. The cultivated cell layer was then plated out on an LB + Kan + Tet plate and incubated overnight to select for trans-conjugates.

#### One-step regime

The donor, a SBW25 strain with a Kan^R^-tagged I55 (with or without *intY*, strains MPB39465/MPB39467 or MPB39565/MPB39568, respectively), and the recipient, a SBW25 strain with a Tet^R^ marker (strain MPB39500), were cultivated separately for 48 h for pre-conditioning. Then 25 μL of each culture and 50 μL of community filtrate from frozen stocks (autoclaved or not) were inoculated into the same microcosm and then cultivated for a further 48 h. A dilution series of the culture was then spread plated on LB + Kan + Tet plates, which were then cultivated for 24 h. The number of colonies were counted and recorded.

#### Two-step regime

For the first step, the donor, a SBW25 strain with a Kan^R^-tagged I55 (or its derivatives), were cultivated for 48 h for pre-conditioning. Then 50 μL of the donor culture, with 5 μL additives (called additive ①) where applicable, was added to microcosms and then cultivated for a further 48 h. A stock of donor-culture filtrate was obtained by filtering the culture through a 0.2 μm filter (Whatman). The filtrate was distributed into 100 μL aliquots. Aliquots were directly frozen in -80 °C for storage. To retrieve the donor-culture filtrate, one aliquot from the frozen stock was kept at room temperature for ∼10 min.

For the second step, the recipient, a SBW25 strain with a Tet^R^ marker (strain MPB39500), was cultivated for 48 h for pre-conditioning. Then 50 μL of the recipient culture, with 5 μL donor culture filtrate (from the stock generated in the first step of the assay, and treated before use where applicable), and with 5 μL additives (called additive ②) where applicable, was added to microcosms and then cultivated for a further 48 h. To test the presence of transfer events of I55, 3 μL of the culture was spotted on LB + Kan + Tet plates, which were then cultivated for 24 h before pictures were taken. To quantify the density of recipient cells that received I55, a dilution series of the culture was spread plated on LB + Kan + Tet plates, which were then cultivated for 24 h before the number of colonies was recorded. To estimate the proportion of recipient cells that received I55, a dilution series of the culture was spread plated on LB + Kan + Tet plates and on LB + Tet plates, which were then cultivated for 24 h before the number of colonies was recorded and divided for proportion.

In the first step, the additive ① was in general community filtrate from the stock of the ‘fixed’ regime, or pure bacteriophage lysates. In cases where the density of agents that were capable of transferring I55 (but not necessarily carrying I55 at this point) in the additive ① was to be estimated, a dilution series of the additive was made and used as separate treatments (called a dilution-to-extinction treatment). In the second step, the additive ② was in general community filtrate from the stock of the ‘fixed’ regime, or pure bacteriophage lysates. In cases where the density of agents that were carrying I55 in the donor culture filtrate or additive ② was to be estimated, a dilution series of the filtrate or additive was made and used as separate treatments (called a dilution-to-extinction treatment). Where applicable, the additives and filtrate samples in first and / or second steps were treated with RNase-Free DNase I (Norgen) or autoclaved before used in the assay.

### Fluorescent fitness assay

The protocol was adapted from Barnett et al. (Barnett et al., 2025). Two competitor strains, one with a GFP tag and the other with an RFP tag, were cultivated separately for 48 h for pre-conditioning. Then 25 μL of each culture, and additives (50 μL of community filtrate or 2 μL of phage lysate, see following sections) from frozen stocks were inoculated into the same microcosm (with 5 mL fresh medium), which was then cultivated for a further 48 h. At the beginning and end of the second incubation period, the ratios of GFP to RFP cells were determined by flow cytometry (MACSQuant VYB), denoted by *r*_0_ and *r*_1_, and total densities of cells in the microcosm were determined by spread plating on LB plates, denoted by *n*_0_ and *n*_1_. The relative fitness of GFP to RFP cells is the ratio of Malthusian parameters (Lenski et al., 1991), given by

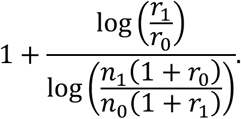

The relative fitness of RFP to GFP cells is the reciprocal of the above expression. To eliminate any fitness difference due to the choice of fluorescent protein markers, the reverse tagging strategy of the two competitor strains with fluorescent protein markers was also implemented to calculate relative fitness.

### Filtrate sequencing and analysis

To sequence each culture filtrate sample, a 100 μL aliquot of filtrate was extracted with Norgen Phage DNA Isolation kit. The concentrations of eluted DNA were in general lower than ∼1 ng/μL and insufficient for subsequent sequencing. To amplify the DNA, 10 μL of eluted DNA was amplified with Cytiva GenomiPhi V3 Ready-To-Go DNA Amplification kit. The amplified DNA was extracted again with Qiagen QIAquick PCR purification kit. The DNA samples Illumina short-read sequenced (read length 150 bp) by Eurofins Scientific SE for ∼15M paired-end reads. Bioinformatic analysis of sequencing data was done in Geneious Prime 2022.2.1 (https://www.geneious.com). For each sample, raw reads were trimmed by BBDuk, and the trimmed reads were also *de novo* assembled by the assembler MEGAHIT (Li et al., 2015)

### Plaque and spot assays of bacteriophages

To determine the phage titer, a dilution series of community filtrate or phage lysate (see following sections) was made. For plaque assays, 50 μL diluted material and 100-400 μL bacterial host overnight culture were mixed into 4 mL SSA (kept at 55 °C). The mixture was immediately poured onto a plate, dried in a laminar flow cabinet for a few minutes, before being incubated for ∼ 36 h. For spot assays, 100-400 μL bacterial host overnight culture were added into 4 mL SSA (kept at 55 °C). The mixture was immediately poured onto a plate and dried in a laminar flow cabinet for a few minutes. Then 3 μL of diluted material was spotted onto the plate and dried in a laminar flow cabinet for another few minutes, before being incubated for ∼ 36 hrs. For both assays, the number of plaques or presence of bacteriophage spots were determined after incubation. Pictures of the plates were taken with Bio-Rad ChemiDoc Imager, under the colorimetric setting with exposure time 0.1 sec.

### Bacteriophage isolation, storage and sequencing

To purify phages from the community filtrate, a plaque assay (in LB environment) using the filtrate was performed. A single isolated plaque was then picked, with which a plaque assay was then performed. This was repeated at least three times. Finally, a single isolated plaque from the last assay was picked and amplified in exponential-phase bacterial host culture in LB medium for ∼6 h. The obtained phage lysate was chloroform extracted and stored in -80 °C.

To purify the JPTI55 jumbo bacteriophage from culture filtrate samples (see Results for details), a plaque assay (in M9-glucose environment) using the filtrate was performed. Single isolated plaques were randomly picked and screened for the JPTI55 genome by PCR. Another plaque assay was then performed with those plaques that were formed by JPTI55. This was repeated at least three times. Finally, a single isolated plaque from the last assay was picked and amplified in exponential-phase bacterial host culture in M9-glucose medium for ∼72 h. The obtained phage lysate was filtered through 0.2 μm filter (Whatman) and stored in -80 °C.

Amplification of all phage isolates was done by inoculating an aliquot of frozen phage lysate into exponential-phase SBW25 culture in M9-glucose medium. After incubation for ∼72 h, the culture was filtered through 0.2 μm filter (Whatman). The filtrate was then aliquoted and directly stored in -80 °C.

For sequencing of the phage isolates, 500 μL frozen phage lysate was extracted by Norgen Phage DNA Isolation kit. The DNA samples were then Illumina short-read sequenced (read length 150 bp) by Eurofins Scientific SE for ∼15M paired-end reads. Bioinformatic analysis of sequencing data was done in Geneious Prime 2022.2.1. For each sample, raw reads were trimmed by BBDuk and mapped to *P. fluorescens* SBW25 reference genome under the default settings. The unmapped reads were saved and then *de novo* assembled into a circular contig. The obtained contig was blasted (Camacho et al., 2009) for similar records in database, from which a phylogeny of its close relatives was built by VICTOR (Meier-Kolthoff & Göker, 2017).

### Statistical analysis

All experiments were done for at least 3 independent biological replicates. Statistical analyses were performed with GraphPad Prism (v. 10.1.2). Welch’s *t*-test and Mann-Whitney test were performed for parametric and non-parametric pairwise comparisons, respectively.

## Supporting information

This document contains all supplementary figures and tables.

## Miscellaneous

### Data availability

Supporting information, including sequencing raw reads, new reference sequences, phage sequences, and bioinformatic codes will be deposited in Edmond (https://edmond.mpg.de/).

### Author contributions

Y.Z., A.D.F., D.W.R. and P.B.R. designed research; Y.Z., Y.M. and C.V. performed research; Y.Z., Y.M., A.D.F., D.W.R. and P.B.R. analyzed data; and Y.Z. and P.B.R. wrote the paper; all authors approved the final version.

### Acknowledgements

We acknowledge the Max Planck Society for generous core funding. We thank members of Department of Microbial Population Biology (past and present) for support, technical assistance and discussion.

