## Supplementary material for "Jumbo phage-mediated transduction of genomic islands": This document contains all supplementary figures and tables.

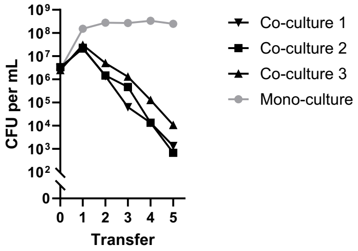


**Fig. E1**. SBW25 density in the co-culture regime (see Fig. 1a(ii) and Materials and Methods) SBW25 co-cultured with compost C community decreased in abundance over transfers. By the 5th transfer the density was only $10^3$ ~ $10^4$ cells (per mL) across the three replicate lines. By contrast, mono-culture of SBW25 maintained a density of $10^8$ ~ $10^9$ cells (per mL).

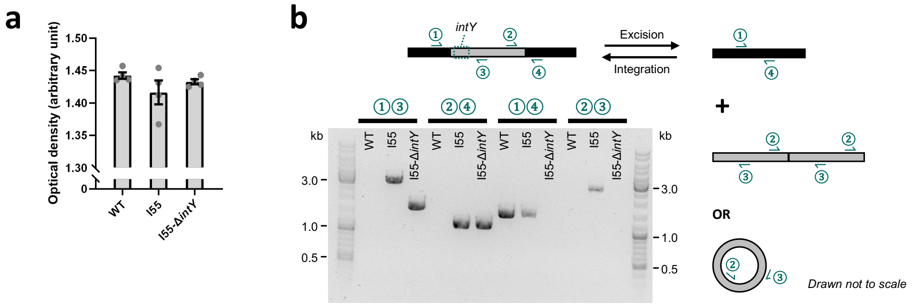


### Fig. E2. Excision and circularization or linear concatenation of I55

**a**. The optical density of overnight LB shaking cultures of SBW25-I55, SBW25-I55Δ*intY* and wild type (WT). Data are mean ± s.d. Brown-Forsythe and Welch ANOVA test was applied to test the difference between the three strains ($p$ = 0.3551).

**b**. PCR of overnight cultures of SBW25-I55, SBW25-I55Δ*intY* and WT, with combination of primers ①②③④. Photos of 3 more biological replicates can be found in Fig. S6. Primers ① and ② flank the left insertion junction, and primers ③ and ④ flanks the right. Primers ① and ④ match the genome, and primers ② and ③ match I55. The presence of bands on SBW25-I55 with pairs ①+② and ③+④ confirms the integration of I55 in the genome. The presence of band on SBW25-I55 with pair ①+④, at a length that is the same as WT, suggests that in the overnight culture of SBW25-I55 there are some cells with I55 excised from the genome. The presence of band on SBW25-I55 with pair ②+③ suggests that I55 circularizes or concatenates so that primers ② and ③ flank the junction. Sanger sequencing of the PCR products of ②+③ confirms that the circular or concatenated I55 contains exactly one 14 bp sequence at the junction, and of ①+④ confirms that I55 excises in a scar-free manner. We therefore proposed that there are two forms of I55 in the population of overnight culture of SBW25-I55, the integrated form (which gives bands with pairs ①+② and ③+④) and the circular form (which gives bands with pairs ①+④ and ②+③). To test whether the tyrosine integrase (IntY) is necessary for excision, we deleted *intY* and did the same PCR. The strain SBW25-I55Δ*intY* showed bands with pairs ①+② and ③+④, but no band with pairs ①+④ and ②+③. This confirms that IntY is necessary for excision and circularization or linear concatenation of I55.

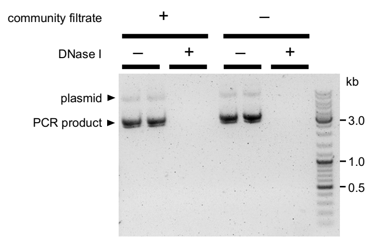


**Fig. E3**. Testing activity of DNase I in community filtrate

To test the efficacy of the RNase-free DNase I (Norgen) in culture filtrate, the filtrate was spiked with linear PCR products and extracted circular plasmids. The mixture was then treated with the DNase according to the manufacturer's instructions, and visualized by gel electrophoresis. Results showed that the DNase I is active in community filtrate.

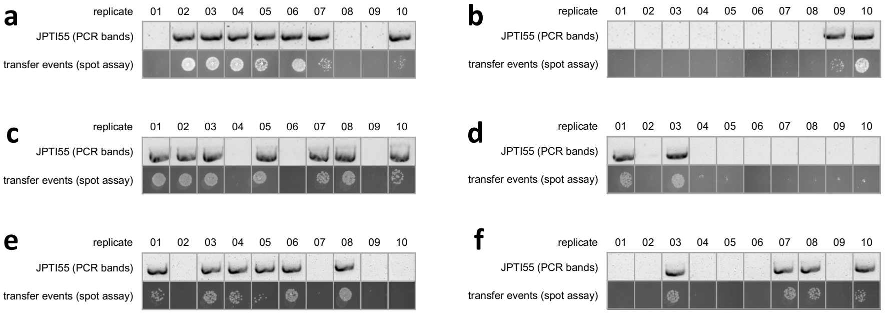


Fig. S1. Full replicates of Fig. 3g.

The example shown in Fig. 3g is panel a here.

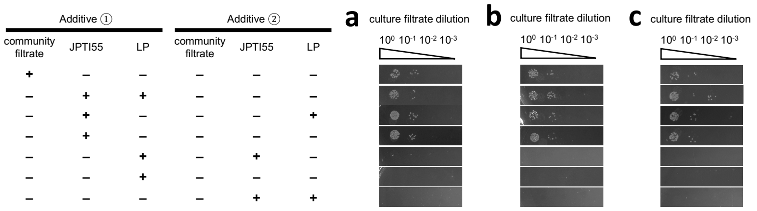


Fig. S2. Full replicates of Fig. 3i.

The example shown in Fig. 3i is panel a here.

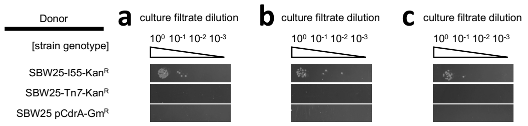


### Fig. S3. Full replicates of Fig. 3j.

The example shown in Fig. 3j is panel a here.

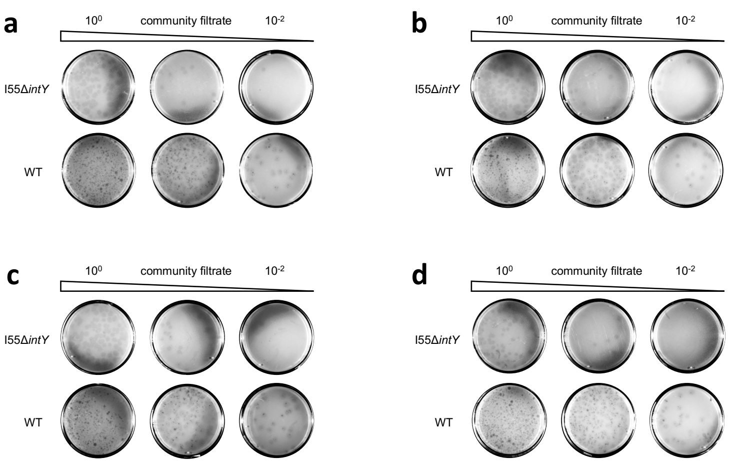


Fig. S4. Full replicates of Fig. 4c.

The example shown in Fig. 4c is panel a here.

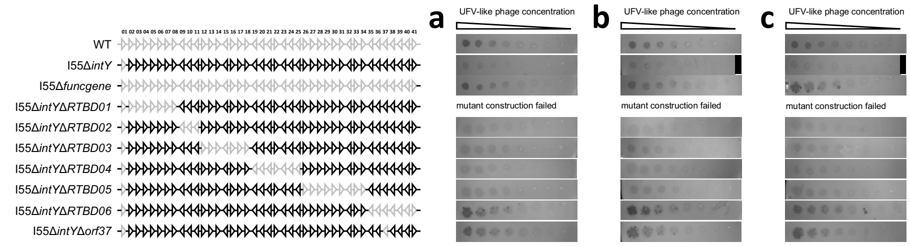


### Fig. S5. Full replicates of Fig. 4f.

The example shown in Fig. 4f is panel a here.

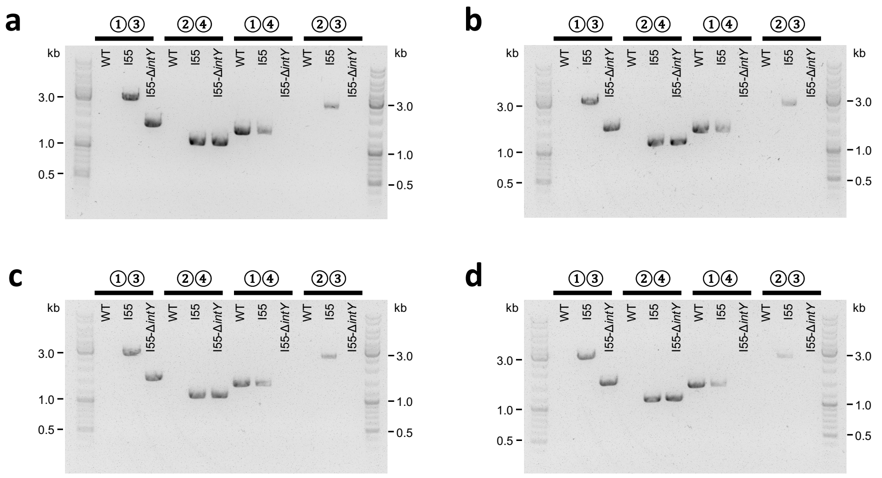


Fig. S6. Full replicates of Fig. E2b.

The example shown in Fig. E2b is panel a here.

### **Table S1**. List of bacterial strains

| Number | Name | Genotype | Construction notes |
|---|---|---|---|
| MPB29212 | SBW25_Wild_Type | SBW25 | Lab stock |
| MPB28806 | SBW25 Tn7-PntpII-mScarlet-I-Gm | SBW25 *glmS*::Tn*7*-145(RFP, Gm$^R$) | Lab stock |
| MPB29284 | Tn7_152_CFC_LB_SBW25 | SBW25 *glmS*::Tn*7*-152(GFP, Kan$^R$) | Conjugation of MPB29212 with MPB27708 |
| MPB29300 | Tn7_155_CFC_LB_SBW25 | SBW25 *glmS*::Tn*7*-155(RFP, Kan$^R$) | Conjugation of MPB29212 with MPB27709 |
| MPB38392 | SFF_b_GFP_4_iso24_23kb | SBW25 *tmRNA*::I23 *glmS*::Tn*7*-152(GFP, Kan$^R$) isolate1* | Isolated from fixed-filtrate experiment, compost b, GFP, line 4 |
| MPB38393 | SFF_b_GFP_5_iso02_23kb | SBW25 *tmRNA*::I23 *glmS*::Tn*7*-152(GFP, Kan$^R$) isolate2* | Isolated from fixed-filtrate experiment, compost b, GFP, line 5 |
| MPB38394 | SFF_b_GFP_5_iso04_23kb | SBW25 *tmRNA*::I23 *glmS*::Tn*7*-152(GFP, Kan$^R$) isolate3* | Isolated from fixed-filtrate experiment, compost b, GFP, line 5 |
| MPB38395 | SFF_b_GFP_5_iso10_23kb | SBW25 *tmRNA*::I23 *glmS*::Tn*7*-152(GFP, Kan$^R$) isolate4* | Isolated from fixed-filtrate experiment, compost b, GFP, line 5 |
| MPB38396 | SFF_b_GFP_5_iso13_23kb | SBW25 *tmRNA*::I23 *glmS*::Tn*7*-152(GFP, Kan$^R$) isolate5* | Isolated from fixed-filtrate experiment, compost b, GFP, line 5 |
| MPB38895 | SFF_b_RFP_3_iso13_23kb | SBW25 *tmRNA*::I23 *glmS*::Tn*7*-155(RFP, Kan$^R$) isolate6* | Isolated from fixed-filtrate experiment, compost b, RFP, line 3 |
| MPB38906 | SFF_b_RFP_4_iso06_44kb | SBW25 *tmRNA*::I44 *glmS*::Tn*7*-155(RFP, Kan$^R$) isolate1* | Isolated from fixed-filtrate experiment, compost b, RFP, line 4 |
| MPB38907 | SFF_b_RFP_4_iso23_44kb | SBW25 *tmRNA*::I44 *glmS*::Tn*7*-155(RFP, Kan$^R$) isolate2* | Isolated from fixed-filtrate experiment, compost b, RFP, line 4 |
| MPB33357 | SF_G_C_2_T14_iso04 | SBW25 *tmRNA*::I55 *glmS*::Tn*7*-152(GFP, Kan$^R$) isolate1* | Isolated from live-filtrate experiment, compost c, GFP, line 2 |
| MPB38103/MPB38109 | Iso04_rev_PFLU4414 | SBW25 *tmRNA*::I55 *glmS*::Tn*7*-152(GFP, Kan$^R$) *pflu3805*SNP | 2SAR of two independent colonies of MPB33357 with MPB38033 |
| MPB38572/MPB38574 | Iso04_rev_PFLU4414_3805 | SBW25 *tmRNA*::I55 *glmS*::Tn*7*-152(GFP, Kan$^R$) | 2SAR of MPB38103/MPB38109 with MPB38391 |
| MPB39024/MPB39026 | Iso04_rev_PFLU4414_3805_del_Tn7 | SBW25 *tmRNA*::I55 | 2SAR of MPB38572/MPB38574 with MPB38943 |
| MPB39132/MPB39136 | SBW25_I55_Δint | SBW25 *tmRNA*::(I55 Δ*intY*) | 2SAR of MPB39024/MPB39026 with MPB38939 |
| MPB39504/MPB39507 | SBW25_I55_Δint_Tn7_GFP | SBW25 *tmRNA*::(I55 Δ*intY*) *glmS*::Tn*7*-152(GFP, Kan$^R$) | Conjugation of MPB39132/MPB39136 with MPB27708 |
| MPB39514/MPB39516 | SBW25_I55_Δint_Tn7_RFP | SBW25 *tmRNA*::(I55 Δ*intY*) *glmS*::Tn*7*-152(GFP, Kan$^R$) | Conjugation of MPB39132/MPB39136 with MPB27709 |
| MPB39379/MPB39459 | SBW25_reconstructed | SBW25 | 2SAR of MPB39132/MPB39136 with MPB39223 |
| MPB39508/MPB39510 | SBW25_reconstructed_Tn7_GFP | SBW25 *glmS*::Tn*7*-152(GFP, Kan$^R$) | Conjugation of MPB39379/MPB39459 with MPB27708 |
| MPB39518/MPB39520 | SBW25_reconstructed_Tn7_RFP | SBW25 *glmS*::Tn*7*-155(RFP, Kan$^R$) | Conjugation of MPB39379/MPB39459 with MPB27709 |
| MPB39465/MPB39467 | SBW25_I55_KanR_InsertS1 | SBW25 *tmRNA*::(I55 (*orf08-orf09*)::Kan$^R$) | 2SAR of MPB39024/MPB39026 with MPB39269 |
| MPB39565/MPB39568 | SBW25_I55_Δint_ KanR_InsertS1 | SBW25 *tmRNA*::(I55 Δ*intY* (*orf08-orf09*)::Kan$^R$) | 2SAR of MPB39132/MPB39136 with MPB39269 |
| MPB39500 | SBW25-TetR | SBW25 *glmS*::Tn7(AGACTGTTGCGCGCTGGCGGTGGCG, Tet$^R$) | Lab stock, see https://doi.org/10.1007/s00239-023-10103-6 |
| MPB16791 | pCdrA-GFP | SBW25 pCdrA-GFP | Lab stock |
| MPB40941 | 55Ele_del_funcgene | SBW25 *tmRNA*::(I55 Δ*funcgene*) ** | 2SAR of MPB39132 with MPB40717 |
| MPB42170 | RTDB_02_del | SBW25 *tmRNA*::(I55 Δ*intY* Δ*RTBD02*) ** | 2SAR of MPB39132 with MPB41414 |
| MPB41585 | RTDB_03_del | SBW25 *tmRNA*::(I55 Δ*intY* Δ*RTBD03*) ** | 2SAR of MPB39132 with MPB41415 |
| MPB42372 | RTDB_04_del | SBW25 *tmRNA*::(I55 Δ*intY* Δ*RTBD04*) ** | 2SAR of MPB39132 with MPB41416 |
| MPB42031 | RTDB_05_del | SBW25 *tmRNA*::(I55 Δ*intY* Δ*RTBD05*) ** | 2SAR of MPB39132 with MPB41417 |

| | | | |
|---|---|---|---|
| MPB41438 | RTDB_06_del | SBW25 *tmRNA*::(I55 Δ*intY* Δ*RTBD06*) ** | 2SAR of MPB39132 with MPB41418 |
| MPB42855 | del_I55orf37_MPB42701 | SBW25 *tmRNA*::(I55 Δ*intY* Δ*orf37*) | 2SAR of MPB39132 with MPB42701 |
| MPB43008 | del_I55orf37_MPB42855_GFP | SBW25 *tmRNA*::(I55 Δ*intY* Δ*orf37*) *glmS*::Tn*7*-152(GFP, Kan$^{R}$) | Conjugation of MPB42855 with MPB27708 |
| MPB43010 | del_I55orf37_MPB42855_Scarlet (FP) | SBW25 *tmRNA*::(I55 Δ*intY* Δ*orf37*) *glmS*::Tn*7*-155(RFP, Kan$^{R}$) | Conjugation of MPB42855 with MPB27709 |

* For variations/mutations, see Table S5.

** Regions funcgene, RTBD01, RTBD02, RTBD03, RTBD04, RTBD05, RTBD06 represents the fragments of I55 between and including the following open reading frames: *orf01* to *orf41*, *orf01* to *orf08*, *orf09* to *orf11*, *orf12* to *orf18*, *orf19* to *orf25*, *orf26* to *orf34*, *orf35* to *orf41*.

**Table S2**. List of *E. coli* plasmids

| Number | Name | Description | Source |
|---|---|---|---|
| MPB27708 | pMRE-Tn7-152 | Plasmid to tag *Pseudomonas* with GFP and Kan$^{R}$ | Gift from Mitja Remus-Emsermann, see https://www.addgene.org/118566/ |
| MPB27709 | pMRE-Tn7-155 | Plasmid to tag *Pseudomonas* with RFP and Kan$^{R}$ | Gift from Mitja Remus-Emsermann, see https://www.addgene.org/118569/ |
| MPB35522 | pUIsacB | Empty plasmid for 2-step allelic replacement | Lab stock, see https://www.biorxiv.org/content/10.1101/2024.07.11.603030v1 |
| MPB35623 | pRK2013 | Helper plasmid for 2-step allelic replacement | Lab stock, see https://www.biorxiv.org/content/10.1101/2024.07.11.603030v1 |
| MPB38033 | rev_PFLU4414 | Plasmid to revert the SNP in *pflu4414* | this study |
| MPB38391 | rev_PFLU3805 | Plasmid to revert the SNP in *pflu3805* | this study |
| MPB38943 | del_Tn7 | Plasmid to delete the Tn*7* cassette | this study |
| MPB38939 | del_55kbInt | Plasmid to delete the gene *intY* of I55 | this study |
| MPB39223 | del_tmIns | Plasmid to delete any inserts downstream of *tmRNA* | this study |
| MPB39269 | KanR_InsertS1 | Plasmid to insert a KanR cassette between *orf08* and *orf09* of I55 | this study |
| MPB41413 | RTBD_01_del | Plasmid to delete region RTBD01 of I55 | this study |
| MPB41414 | RTBD_02_del | Plasmid to delete region RTBD02 of I55 | this study |
| MPB41415 | RTBD_03_del | Plasmid to delete region RTBD03 of I55 | this study |
| MPB41416 | RTBD_04_del | Plasmid to delete region RTBD04 of I55 | this study |
| MPB41417 | RTBD_05_del | Plasmid to delete region RTBD05 of I55 | this study |
| MPB41418 | RTBD_06_del | Plasmid to delete region RTBD06 of I55 | this study |
| MPB42701 | del_I55orf37 | Plasmid to delete *orf37* of I55 | this study |

### **Table S3**. List of primers

| **Number** | **Name** | **Purpose** | **Sequence** |
|---|---|---|---|
| YZ001 | YZ_SBW_5797955F | To detect potential insertions downstream of tmRNA | GTTACCGCTGCGATGTTGAC |
| YZ002 | YZ_SBW_5799313R | To detect potential insertions downstream of tmRNA | GCGTGCTGATGCTGCTAATC |
| YZ003 | YZ_In_1016R | To confirm the presence of I55 | GTGGCCAGATCAGGGAAGAC |
| YZ004 | YZ_In_54359F | To confirm the presence of I55 | ACACGACACACGAAGAGACC |
| YZ005 | pUIsacB_fwd | To amplify the pUIsacB plasmid backbone | AGCACTACATCAACTGACTA |
| YZ006 | pUIsacB_rev | To amplify the pUIsacB plasmid backbone | CTGAACCAAGATAGCTGTAC |
| YZ007 | 2SAR_4880278_Insert_fwd | To construct the plasmid for reverting the SNP in *pflu4414* | GTACAGCTATCTTGGTTCAGGAAATCAACCCATCGCCTG |
| YZ008 | 2SAR_4880278_Insert_rev | To construct the plasmid for reverting the SNP in *pflu4414* | TAGTCAGTTGATGTAGTGCTAACGCCTGTCCTATGTGG |
| YZ009 | pUIsacB_seq_fwd | To Sanger sequence the assembled regions of the pUIsacB plasmid | ATTTGTCCTACTCAGGAGAGCGTTC |
| YZ010 | 4880278_Insert_4962R | To Sanger sequence the plasmid for reverting the SNP in *pflu4414* | GCTGAGTTCGAATCCCTGCT |
| YZ011 | 4880278_Insert_4804F | To Sanger sequence the plasmid for reverting the SNP in *pflu4414* | CTTGCCATGCAGTTGGTCG |
| YZ012 | 4880278_Insert_5486R | To Sanger sequence the plasmid for reverting the SNP in *pflu4414* | GGCCGGCGAAATTTTAGAGC |
| YZ013 | 4880278_Insert_5275F | To Sanger sequence the plasmid for reverting the SNP in *pflu4414* | CGAATCAACGTGACGCTCAC |
| YZ014 | 4880278_Insert_5956R | To Sanger sequence the plasmid for reverting the SNP in *pflu4414* | CGACTGGGGCCGTTTCAT |
| YZ015 | 4880278_Insert_5777F | To Sanger sequence the plasmid for reverting the SNP in *pflu4414* | ACACGCTTGATCACCTGACC |
| YZ016 | pUIsacB_seq_rev | To Sanger sequence the assembled regions of the pUIsacB plasmid | AGCGTTCTGAACAAATCCAGATG |
| YZ084 | SBW_23kb_insert_1 | To confirm the presence of I23 | CCGTTTCTCTTTGGGGTCGA |
| YZ085 | SBW_23kb_insert_2 | To confirm the presence of I23 | ACTCAAGAGCCTCGCCAAAG |
| YZ086 | 2SAR_rev_3805_1 | To construct the plasmid for reverting the SNP in *pflu3805* | GTACAGCTATCTTGGTTCAGCCAGCAACGGGTTGTTCTTG |
| YZ087 | 2SAR_rev_3805_2 | To construct the plasmid for reverting the SNP in *pflu3805* | TAGTCAGTTGATGTAGTGCTCCACTGATCTTGACGCCAGA |
| YZ088 | rev_PFLU3805_4997F | To Sanger sequence the plasmid for reverting the SNP in *pflu3805* | CCTGTTCGCTGAAAATGGCC |
| YZ089 | rev_PFLU3805_5687R | To Sanger sequence the plasmid for reverting the SNP in *pflu3805* | ACACGATGACGACGGTTCTG |
| YZ091 | PFLU4414_Final_Seq_2 | To confirm the genotype of mutants after allelic replacement in *pflu4414* | GAACGCATCCACGGCTAATG |
| YZ092 | PFLU3805_Final_Seq_1 | To confirm the genotype of mutants after allelic replacement in *pflu3805* | GACACGTCGACCTTCTGGAAG |
| YZ093 | PFLU3805_Final_Seq_2 | To confirm the genotype of mutants after allelic replacement in *pflu3805* | CGAGTTGATTGGGGTGGCC |
| YZ098 | PFLU4414_Final_Seq_1_1 | To confirm the genotype of mutants after allelic replacement in *pflu4414* | CGCCGACCATGTTCATGATG |
| YZ102 | SBW_44kb_insert_1 | To confirm the presence of I44 | GCATAGGCTGTGGTCACTGA |
| YZ103 | SBW_44kb_insert_2 | To confirm the presence of I44 | AGACCAAGGCCGTCAAGAAG |
| YZ105 | 2SAR_del_55kbInt_1 | To construct the plasmid for deleting the tyrosine integrase (*orf01*) of I55 | GTACAGCTATCTTGGTTCAGGTTTAGGATTCGACGCCG |
| YZ106 | 2SAR_del_55kbInt_2 | To construct the plasmid for deleting the tyrosine integrase (*orf01*) of I55 | GAAGTCTCTCGGGTGTATTCCAGTGCTGTTTATG |
| YZ107 | 2SAR_del_55kbInt_3 | To construct the plasmid for deleting the tyrosine integrase (*orf01*) of I55 | TGGAATACACCCGAGAGACTTCAACCAGTGA |

| | | | |
|---|---|---|---|
| YZ108 | 2SAR_del_55kbInt_4 | To construct the plasmid for deleting the tyrosine integrase (*orf01*) of I55 | TAGTCAGTTGATGTAGTGCTTCTTCGTTCACCACATTCCG |
| YZ109 | del_55kbInt_4941F | To Sanger sequence the plasmid for deleting the tyrosine integrase (*orf01*) of I55 | TGGACGAAGCGGCTAAAACT |
| YZ110 | del_55kbInt_5514R | To Sanger sequence the plasmid for deleting the tyrosine integrase (*orf01*) of I55 | TCGCCATTCAGCTGTCCAAT |
| YZ111 | del_55kbInt_Final_Seq_1 | To confirm the genotype of mutants after allelic re placement in the tyrosine integrase (*orf01*) of I55 | ACGTTAGCTCAGCCGAGGT |
| YZ112 | 2SAR_del_Tn7_1 | To construct the plasmid for deleting the Tn*7* cassette | GTACAGCTATCTTGGTTCAGTGAGCTATCAGTTGCGCCTG |
| YZ113 | 2SAR_del_Tn7_2 | To construct the plasmid for deleting the Tn*7* cassette | TAGTCAGTTGATGTAGTGCTAATCCGACCTGACCTTGCTG |
| YZ114 | del_Tn7_4864F | To Sanger sequence the plasmid for deleting the Tn*7* cassette | TTGCGGGTTACGGATGACC |
| YZ115 | del_Tn7_5273F | To Sanger sequence the plasmid for deleting the Tn*7* cassette | GACGTTGGTGGATCAGGGAG |
| YZ116 | del_Tn7_5670F | To Sanger sequence the plasmid for deleting the Tn*7* cassette | CAATGGGCCGTGTTTGAGC |
| YZ117 | del_Tn7_5084R | To Sanger sequence the plasmid for deleting the Tn*7* cassette | TTGGAAGGTGAGCTGCTGTC |
| YZ118 | del_Tn7_5480R | To Sanger sequence the plasmid for deleting the Tn*7* cassette | CTGTCGCCGATCCTCTACAC |
| YZ119 | del_Tn7_5845R | To Sanger sequence the plasmid for deleting the Tn*7* cassette | CAATGGACAGCACCGTGGAA |
| YZ120 | del_Tn7_Final_Seq_1 | To confirm the genotype of mutants after allelic replacement in the Tn*7* cassette | ACACCGCCCAACGTGATG |
| YZ121 | del_Tn7_Final_Seq_2 | To confirm the genotype of mutants after allelic replacement in the Tn*7* cassette | ACACCCTGTTCGTGACCATC |
| YZ124 | 2SAR_KanR_InsertS1_1 | To construct the plasmid for inserting an Kan$^R$ cassette between *orf08* and *orf09* of I55 | GTACAGCTATCTTGGTTCAGGTCCGCAGAACTGTTGGAG |
| YZ125 | 2SAR_KanR_InsertS1_2 | To construct the plasmid for inserting an Kan$^R$ cassette between *orf08* and *orf09* of I55 | GATTAGCTGTGTCGCTACTG |
| YZ126 | 2SAR_KanR_InsertS1_3 | To construct the plasmid for inserting an Kan$^R$ cassette between *orf08* and *orf09* of I55 | CCGCTCACCGAAGCTGATTA |
| YZ127 | 2SAR_KanR_InsertS1_4 | To construct the plasmid for inserting an Kan$^R$ cassette between *orf08* and *orf09* of I55 | TAGTCAGTTGATGTAGTGCTGGCGACTTCGGTATTCCAG |
| YZ128 | 2SAR_KanR_InsertS1_5 | To construct the plasmid for inserting an Kan$^R$ cassette between *orf08* and *orf09* of I55 | CAGTAGCGACACAGCTAATCGGGAGACCAGAAACAAAAAAACACCC CGAAGGGTGTTCAATAATTGGTCCGCTTCCTTTAGCAGCC |
| YZ129 | 2SAR_KanR_InsertS1_6 | To construct the plasmid for inserting an Kan$^R$ cassette between *orf08* and *orf09* of I55 | TAATCAGCTTCGGTGAGCGGTCCTAGGTGCTCGAGTGG |
| YZ130 | KanR_InsertS1_5008F | To Sanger sequence the plasmid for inserting an Kan$^R$ cassette between *orf08* and *orf09* of I55 | TGGCACACGATACCGAAGAC |
| YZ131 | KanR_InsertS1_5508F | To Sanger sequence the plasmid for inserting an Kan$^R$ cassette between *orf08* and *orf09* of I55 | AGCGTGCGAGCTCTAAAACA |
| YZ132 | KanR_InsertS1_6011F | To Sanger sequence the plasmid for inserting an Kan$^R$ cassette between *orf08* and *orf09* of I55 | ACTCTTCCGAGCAAAGGACG |
| YZ133 | KanR_InsertS1_6662F | To Sanger sequence the plasmid for inserting an Kan$^R$ cassette between *orf08* and *orf09* of I55 | GCAGGAAATGGGTCATCGGT |
| YZ134 | KanR_InsertS1_5189R | To Sanger sequence the plasmid for inserting an Kan$^R$ cassette between *orf08* and *orf09* of I55 | TTGCAGCAGTGACAAGGACA |
| YZ135 | KanR_InsertS1_5695R | To Sanger sequence the plasmid for inserting an Kan$^R$ cassette between *orf08* and *orf09* of I55 | TTGATCTTGGGAGAAGCGGC |
| YZ136 | KanR_InsertS1_6177R | To Sanger sequence the plasmid for inserting an Kan$^R$ cassette between *orf08* and *orf09* of I55 | GGACAGCCGGTATAAAGGGA |
| YZ137 | KanR_InsertS1_6853R | To Sanger sequence the plasmid for inserting an Kan$^R$ cassette between *orf08* and *orf09* of I55 | TCTCACACGTTCGTTCGACC |
| YZ144 | 2SAR_del_tmIns_1 | To construct the plasmid for deleting MGEs inserted downstream of *tmRNA* | GTACAGCTATCTTGGTTCAGGAGATCGACACCGCCGTCTC |
| YZ145 | 2SAR_del_tmIns_2 | To construct the plasmid for deleting MGEs inserted downstream of *tmRNA* | TAGTCAGTTGATGTAGTGCTGCGTGCTGATGCTGCTAATC |
| YZ146 | del_tmIns_5225F | To Sanger sequence the plasmid for deleting MGEs inserted downstream of *tmRNA* | GGGTCGCTGAGGGTTAACTT |
| YZ147 | del_tmIns_5699F | To Sanger sequence the plasmid for deleting MGEs inserted downstream of *tmRNA* | ATGACGCTCGTGTCCAACTT |
| YZ148 | del_tmIns_4886R | To Sanger sequence the plasmid for deleting MGEs inserted downstream of *tmRNA* | CTGTGCAGTGTACGCCAAAC |
| YZ149 | del_tmIns_5575R | To Sanger sequence the plasmid for deleting MGEs inserted downstream of *tmRNA* | GGTACAGACCCTGTTCGTCG |

| | | | |
|---|---|---|---|
| YZ150 | del_tmIns_Final_Seq_1 | To confirm the genotype of mutants after allelic replacement downstream of *tmRNA* | TTACCGCTGCGATGTTGACT |
| YZ151 | del_tmIns_Final_Seq_2 | To confirm the genotype of mutants after allelic replacement downstream of *tmRNA* | GGAGCGTCCGTGAGAGATTC |
| YZ161 | Tn7R-109 | To confirm the insertion of Tn7 cassette in SBW25 genome | CAGCATAACTGGACTGATTTCAG |
| YZ162 | SBW25-glmS | To confirm the insertion of Tn7 cassette in SBW25 genome | CACCAAAGCTTTCACCACCCAA |
| YZ242 | Jumbo_Region1_fwd | To screen the presence of the jumbo phage JPTI55 | GAAGTCGTACTGGGCACACA |
| YZ243 | Jumbo_Region1_rev | To screen the presence of the jumbo phage JPTI55 | GTCACACCACGCACTTCAAC |
| YZ248 | UFV_like_1_fwd | To screen the presence of the UFV-like phage | GACCGTCAGGCATAGGTAGC |
| YZ249 | UFV_like_1_rev | To screen the presence of the UFV-like phage | TTCAGGTCGCGCTCTTTCTT |
| CV11 | RTBD_01_upstream_fwd | To construct the plasmid for deleting RTBD_01 of I55 | GTACAGCTATCTTGGTTCAGTAACGTCGCTCCGTCTAAG |
| CV12 | RTBD_01_upstream_rev | To construct the plasmid for deleting RTBD_01 of I55 | CTCGAAACCTGGATTGTTATCTTTCTCAGGTTG |
| CV13 | RTBD_01_downstream_fwd | To construct the plasmid for deleting RTBD_01 of I55 | ATAACAATCCAGGTTTCGAGAGTAGAAAG |
| CV14 | RTBD_01_downstream_rev | To construct the plasmid for deleting RTBD_01 of I55 | TAGTCAGTTGATGTAGTGCTCGTCACACCGAGTTTAAC |
| CV17 | RTBD_02_upstream_fwd | To construct the plasmid for deleting RTBD_02 of I55 | GTACAGCTATCTTGGTTCAGTACCGGCAGCTTCGCCTG |
| CV18 | RTBD_02_upstream_rev | To construct the plasmid for deleting RTBD_02 of I55 | GTGTCCGAGTATCTTCTCGGCAATCCTTCAGG |
| CV19 | RTBD_02_downstream_fwd | To construct the plasmid for deleting RTBD_02 of I55 | CCGAGAAGATACTCGGACACCTCGGCGG |
| CV20 | RTBD_02_downstream_rev | To construct the plasmid for deleting RTBD_02 of I55 | TAGTCAGTTGATGTAGTGCTTCGTGAAGTCAGCCGAATTGACG |
| CV23 | RTBD_03_upstream_fwd | To construct the plasmid for deleting RTBD_03 of I55 | GTACAGCTATCTTGGTTCAGCTCAAAGCGAAGCCAATAC |
| CV24 | RTBD_03_upstream_rev | To construct the plasmid for deleting RTBD_03 of I55 | GGTGAACGGCGGGGATATCTCTCACTGG |
| CV25 | RTBD_03_downstream_fwd | To construct the plasmid for deleting RTBD_03 of I55 | AGATATCCCCGCCGTTCACCGAGCACC |
| CV26 | RTBD_03_downstream_rev | To construct the plasmid for deleting RTBD_03 of I55 | TAGTCAGTTGATGTAGTGCTCTGACGGTCATAACTCAGAATGTCG |
| CV29 | RTBD_04_upstream_fwd | To construct the plasmid for deleting RTBD_04 of I55 | GTACAGCTATCTTGGTTCAGACTCAAGGAGTGGTCCTCG |
| CV30 | RTBD_04_upstream_rev | To construct the plasmid for deleting RTBD_04 of I55 | GGAAAGCGTTCCTGCAGCGAGAATCGGTC |
| CV31 | RTBD_04_downstream_fwd | To construct the plasmid for deleting RTBD_04 of I55 | TCGCTGCAGGAACGCTTTCCTCAGGTCTG |
| CV32 | RTBD_04_downstream_rev | To construct the plasmid for deleting RTBD_04 of I55 | TAGTCAGTTGATGTAGTGCTCAACTTTTCGCCAAGCTG |
| CV35 | RTBD_05_upstream_fwd | To construct the plasmid for deleting RTBD_05 of I55 | GTACAGCTATCTTGGTTCAGGGTGTCGTCTCAGCCCCC |
| CV36 | RTBD_05_upstream_rev | To construct the plasmid for deleting RTBD_05 of I55 | GTCAGGGTCTGGGGCGAAGCTCCTTGGT |
| CV37 | RTBD_05_downstream_fwd | To construct the plasmid for deleting RTBD_05 of I55 | GCTTCGCCCCAGACCCTGACCACCTAGTG |
| CV38 | RTBD_05_downstream_rev | To construct the plasmid for deleting RTBD_05 of I55 | TAGTCAGTTGATGTAGTGCT ACAAGATCGGTGTTACGG |
| CV41 | RTBD_06_upstream_fwd | To construct the plasmid for deleting RTBD_06 of I55 | GTACAGCTATCTTGGTTCAGGAAGTTCTACCTGGATGCAG |
| CV42 | RTBD_06_upstream_rev | To construct the plasmid for deleting RTBD_06 of I55 | CCGAAGCGCTGTCTGATCGGCTTCGTCTC |
| CV43 | RTBD_06_downstream_fwd | To construct the plasmid for deleting RTBD_06 of I55 | CCGATCAGACAGCGCTTCGGTTTGACTAAAC |
| CV44 | RTBD_06_downstream_rev | To construct the plasmid for deleting RTBD_06 of I55 | TAGTCAGTTGATGTAGTGCTCGAGCGTCATCACGTTTG |
| CV75 | 2SAR_del_I55orf37_1 | To construct the plasmid for deleting *orf37* of I55 | GTACAGCTATCTTGGTTCAGGTCGCGAGCAATGAAGTG |
| CV76 | 2SAR_del_I55orf37_2 | To construct the plasmid for deleting *orf37* of I55 | AATAGCCTCACCTGACTTGCTCCTTTACAGG |
| CV77 | 2SAR_del_I55orf37_3 | To construct the plasmid for deleting *orf37* of I55 | GGAGCAAGTCAGGTGAGGCTATTCCGGTCG |

| | | | |
|---|---|---|---|
| CV78 | 2SAR_del_I55orf37_4 | To construct the plasmid for deleting *orf37* of I55 | TAGTCAGTTGATGTAGTGCTCAAGGGGATCGGAGTGAAG |

### Table S4. Annotations of MGEs I23, I44 and I55

| Element | ORF name | Consensus annotation | Minimum coordinate | Maximum coordinate | Length | Direction of transcription | Prokka product | Pfam domains | DefenseFinder prediction |
|---|---|---|---|---|---|---|---|---|---|
| I23 | *orf01 (intY)* | tyrosine integrase | 186 | 1442 | 1257 | forward | Prophage integrase IntA | PF13356 (Arm-DNA-bind_3; aa 8-94); PF22022 (Phage_int_M; aa 107-200); PF00589 (Phage_integrase; aa 214-378). | NA |
| I23 | *orf02* | GajA | 1631 | 3436 | 1806 | reverse | ATP-dependent DNA helicase Rep | PF00580 (UvrD-helicase; aa 5-125, 133-239). | Gabija__GajB_1. |
| I23 | *orf03* | GajB | 3462 | 5312 | 1851 | reverse | hypothetical protein | PF13175 (AAA_15; aa 1-72, 310-381); PF20469 (OLD-like_TOPRIM; aa 429-500). | Gabija__GajA. |
| I23 | *orf04* | unannotated | 5637 | 5966 | 330 | forward | hypothetical protein | NA | NA |
| I23 | *orf05* | DUF932 | 6062 | 7030 | 969 | forward | hypothetical protein | PF06067 (DUF932; aa 77-302). | NA |
| I23 | *orf06* | YqaJ | 7106 | 8110 | 1005 | forward | hypothetical protein | PF09588 (YqaJ; aa 29-172). | NA |
| I23 | *orf07* | Gp3-like | 8201 | 9151 | 951 | forward | hypothetical protein | PF18897 (Gp3-like; aa 12-206). | NA |
| I23 | *orf08* | RadC | 9123 | 9620 | 498 | forward | hypothetical protein | PF04002 (RadC; aa 44-163). | NA |
| I23 | *orf09* | unannotated | 9634 | 9843 | 210 | forward | hypothetical protein | NA | NA |
| I23 | *orf10* | unannotated | 9903 | 10208 | 306 | forward | hypothetical protein | NA | NA |
| I23 | *orf11* | HTH-containing protein | 10195 | 10542 | 348 | reverse | hypothetical protein | PF01381 (HTH_3; aa 17-71). | NA |
| I23 | *orf12* | unannotated | 10681 | 11034 | 354 | forward | hypothetical protein | NA | NA |
| I23 | *orf13* | unannotated | 11155 | 11973 | 819 | forward | hypothetical protein | NA | NA |
| I23 | *orf14* | DNA topoisomerase | 12150 | 12920 | 771 | reverse | hypothetical protein | PF08378 (NERD; aa 33-145); PF01396 (zf-C4_Topoisom; aa 217-245). | NA |
| I23 | *orf15* | T5orf172 helicase | 13022 | 14206 | 1185 | reverse | hypothetical protein | PF10544 (T5orf172; aa 263-374). | NA |
| I23 | *orf16* | DEAD-like helicase | 14199 | 16265 | 2067 | reverse | hypothetical protein | PF00270 (DEAD; aa 28-193). | NA |
| I23 | *orf17* | unannotated | 16268 | 16372 | 105 | reverse | hypothetical protein | NA | NA |
| I23 | *orf18* | DNA methyltransferase | 16369 | 19131 | 2763 | reverse | hypothetical protein | PF20464 (MmeI_N; aa 1-159); PF20465 (MmeI_hel; aa 170-250); PF20473 (MmeI_Mtase; aa 325-606); PF20466 (MmeI_TRD; aa 629-832); PF20467 (MmeI_C; aa 834-911). | RM_Type_IIG__Type_IIG. |
| I23 | *orf19* | inovirus Gp2 | 19299 | 20000 | 702 | reverse | hypothetical protein | PF11726 (Inovirus_Gp2; aa 50-231). | NA |
| I23 | *orf20 (alpA)* | AlpA | 20507 | 20740 | 234 | reverse | hypothetical protein | PF05930 (Phage_AlpA; aa 1-51). | NA |
| I23 | *orf21* | unannotated | 20810 | 22114 | 1305 | reverse | hypothetical protein | NA | NA |

| | | | | | | | | | |
|---|---|---|---|---|---|---|---|---|---|
| I23 | *orf22* | unannotated | 22346 | 23020 | 675 | forward | hypothetical protein | NA | NA |
| I44 | *orf01 (intY)* | tyrosine integrase | 185 | 1441 | 1257 | forward | Prophage integrase IntA | PF13356 (Arm-DNA-bind_3; aa 8-94); PF22022 (Phage_int_M; aa 107-200); PF00589 (Phage_integrase; aa 214-377). | NA |
| I44 | *orf02* | type II REase | 1609 | 2364 | 756 | forward | hypothetical protein | PF01844 (HNH; aa 180-235). | RM_Type_II__Type_II_REases |
| I44 | *orf03* | type II Mtase | 2467 | 3531 | 1065 | forward | putative BsuMI modification methylase subunit YdiO | PF00145 (DNA_methylase; aa 4-336). | RM_Type_II__Type_II_MTases |
| I44 | *orf04* | VSR endonuclease | 3482 | 3898 | 417 | reverse | Very short patch repair protein | PF03852 (Vsr; aa 2-64). | NA |
| I44 | *orf05* | DrmC | 3916 | 4554 | 639 | reverse | Cardiolipin synthase B | PF13091 (PLDc_2; aa 67-200). | DISARM__drmC |
| I44 | *orf06* | DrmB | 4682 | 6538 | 1857 | reverse | hypothetical protein | PF09369 (MZB; aa 483-583). | DISARM__drmB |
| I44 | *orf07* | DrmA | 6541 | 10575 | 4035 | reverse | hypothetical protein | NA | DISARM__drmA |
| I44 | *orf08* | unannotated | 10579 | 10908 | 330 | reverse | hypothetical protein | NA | NA |
| I44 | *orf09* | DrmMI | 10905 | 14981 | 4077 | reverse | hypothetical protein | PF07669 (Eco57I; aa 845-931). | DISARM_1__drmMI |
| I44 | *orf10* | DrmD | 14978 | 18103 | 3126 | reverse | RNA polymerase-associated protein RapA | PF00176 (SNF2-rel_dom; aa 134-317); PF00271 (Helicase_C; aa 514-620). | DISARM_1__drmD |
| I44 | *orf11* | MFS transporter | 18695 | 19495 | 801 | reverse | Glycerol-3-phosphate transporter | PF07690 (MFS_1; aa 29-257). | NA |
| I44 | *orf12* | alcohol dehydrogenase | 19609 | 20700 | 1092 | reverse | NAD-dependent methanol dehydrogenase | PF00465 (Fe-ADH; aa 8-351). | NA |
| I44 | *orf13* | avidin | 20847 | 21623 | 777 | reverse | hypothetical protein | PF01382 (Avidin; aa 3-85). | NA |
| I44 | *orf14* | MFS transporter | 21691 | 23055 | 1365 | reverse | Putative niacin/nicotinamide transporter NaiP | PF07690 (MFS_1; aa 46-378). | NA |
| I44 | *orf15* | Asp/Glu/hydantoin racemase | 23118 | 23939 | 822 | reverse | Hydantoin racemase | PF01177 (Asp_Glu_race; aa 20-225). | NA |
| I44 | *orf16* | unannotated | 23960 | 24850 | 891 | reverse | hypothetical protein | NA | NA |
| I44 | *orf17* | metallopeptidase | 24907 | 26292 | 1386 | reverse | Xaa-Pro aminopeptidase | PF05195 (AMP_N; aa 7-123); PF00557 (Peptidase_M24; aa 173-438). | NA |
| I44 | *orf18* | hydantoinase B | 26320 | 27978 | 1659 | reverse | Acetophenone carboxylase delta subunit | PF02538 (Hydantoinase_B; aa 5-522). | NA |

| | | | | | | | | | |
|---|---|---|---|---|---|---|---|---|---|
| I44 | *orf19* | hydantoinase A | 27992 | 30076 | 2085 | reverse | Acetophenone carboxylase gamma subunit | PF05378 (Hydant_A_N; aa 9-185); PF01968 (Hydantoinase_A; aa 208-500); PF19278 (Hydant_A_C; aa 514-689). | NA |
| I44 | *orf20* | HTH-type transcriptional regulator | 30253 | 32856 | 2604 | reverse | HTH-type transcriptional regulator MalT | PF00196 (GerE; aa 802-856). | NA |
| I44 | *orf21* | unannotated | 33248 | 33571 | 324 | forward | hypothetical protein | NA | NA |
| I44 | *orf22* | DUF932 | 33664 | 34632 | 969 | forward | hypothetical protein | PF06067 (DUF932; aa 77-302). | NA |
| I44 | *orf23* | YqaJ | 34707 | 35711 | 1005 | forward | hypothetical protein | PF09588 (YqaJ; aa 29-172). | NA |
| I44 | *orf24* | Gp3-like | 35790 | 36740 | 951 | forward | hypothetical protein | PF18897 (Gp3-like; aa 12-206). | NA |
| I44 | *orf25* | RadC | 36712 | 37209 | 498 | forward | hypothetical protein | PF04002 (RadC; aa 44-163). | NA |
| I44 | *orf26* | unannotated | 37223 | 37432 | 210 | forward | hypothetical protein | NA | NA |
| I44 | *orf27* | unannotated | 37534 | 37779 | 246 | forward | hypothetical protein | NA | NA |
| I44 | *orf28* | HTH-containing protein | 37787 | 38134 | 348 | reverse | hypothetical protein | PF01381 (HTH_3; aa 17-71). | NA |
| I44 | *orf29* | unannotated | 38272 | 38658 | 387 | forward | hypothetical protein | NA | NA |
| I44 | *orf30* | unannotated | 38675 | 39493 | 819 | forward | hypothetical protein | NA | NA |
| I44 | *orf31* | inovirus Gp2 | 39957 | 40916 | 960 | reverse | hypothetical protein | PF11726 (Inovirus_Gp2; aa 149-232). | NA |
| I44 | *orf32 (alpA)* | AlpA | 41252 | 41482 | 231 | reverse | hypothetical protein | PF05930 (Phage_AlpA; aa 1-51). | NA |
| I44 | *orf33* | unannotated | 41531 | 42988 | 1458 | reverse | hypothetical protein | NA | NA |
| I44 | *orf34* | unannotated | 43158 | 43829 | 672 | forward | hypothetical protein | NA | NA |
| I55 | *orf01 (intY)* | tyrosine integrase | 185 | 1441 | 1257 | forward | Prophage integrase IntA | PF13356 (Arm-DNA-bind_3; aa 8-94); PF22022 (Phage_int_M; aa 107-200); PF00589 (Phage_integrase; aa 214-378). | NA |
| I55 | *orf02* | dynamin | 2594 | 4903 | 2310 | forward | hypothetical protein | PF00350 (Dynamin_N; aa 59-226). | Eleos__LeoA |
| I55 | *orf03* | dynamin | 4914 | 7166 | 2253 | forward | hypothetical protein | PF00350 (Dynamin_N; aa 40-177). | NA |
| I55 | *orf04* | unannotated | 7235 | 7738 | 504 | forward | hypothetical protein | NA | NA |
| I55 | *orf05* | unannotated | 7735 | 8307 | 573 | forward | hypothetical protein | NA | NA |
| I55 | *orf06* | unannotated | 8349 | 10991 | 2643 | forward | hypothetical protein | NA | NA |
| I55 | *orf07* | unannotated | 11420 | 12151 | 732 | forward | hypothetical protein | NA | NA |
| I55 | *orf08* | unannotated | 12148 | 13686 | 1539 | forward | hypothetical protein | NA | NA |
| I55 | *orf09* | helicase | 14030 | 16783 | 2754 | reverse | RNA polymerase-associated protein RapA | PF04851 (ResIII; aa 36-245); PF00271 (Helicase_C; aa 755-814). | NA |
| I55 | *orf10* | unannotated | 16770 | 18677 | 1908 | reverse | hypothetical protein | NA | NA |
| I55 | *orf11* | unannotated | 18674 | 20056 | 1383 | reverse | hypothetical protein | NA | NA |
| I55 | *orf12* | unannotated | 20681 | 21103 | 423 | forward | hypothetical protein | NA | NA |

| | | | | | | | | | |
|---|---|---|---|---|---|---|---|---|---|
| I55 | *orf13* | DUF1232 | 21186 | 21524 | 339 | forward | hypothetical protein | PF06803 (DUF1232; aa 50-84). | NA |
| I55 | *orf14* | unannotated | 21652 | 22302 | 651 | forward | hypothetical protein | NA | NA |
| I55 | *orf15* | unannotated | 22316 | 22447 | 132 | reverse | hypothetical protein | NA | NA |
| I55 | *orf16* | AAA | 22605 | 24695 | 2091 | forward | ATP-dependent zinc metalloprotease FtsH | PF00004 (AAA; aa 255-377, 491-614). | NA |
| I55 | *orf17* | WYL | 24791 | 25813 | 1023 | forward | hypothetical protein | PF13280 (WYL; aa 156-330). | WYL_I_II_III_IV_V_VI_4 |
| I55 | *orf18* | unannotated | 25859 | 26395 | 537 | forward | hypothetical protein | NA | NA |
| I55 | *orf19* | Sir2 | 26521 | 27174 | 654 | reverse | NAD-dependent protein deacylase | PF02146 (SIR2; aa 20-196). | NA |
| I55 | *orf20* | unannotated | 27411 | 27668 | 258 | reverse | hypothetical protein | NA | NA |
| I55 | *orf21* | Sel1 | 27658 | 28299 | 642 | reverse | hypothetical protein | PF08238 (Sel1; aa 96-127, 133-166, 171-200). | NA |
| I55 | *orf22* | unannotated | 28421 | 28585 | 165 | forward | hypothetical protein | NA | NA |
| I55 | *orf23* | RecD | 28573 | 30666 | 2094 | reverse | RecBCD enzyme subunit RecD | PF21185 (RecD_N; aa 40-153); PF13245 (AAA_19; aa 205-357); PF13538 (UvrD_C_2; aa 615-662). | NA |
| I55 | *orf24* | RecB | 30663 | 34337 | 3675 | reverse | RecBCD enzyme subunit RecB | PF00580 (UvrD-helicase; aa 13-452); PF13361 (UvrD_C; aa 459-660); PF13361 (UvrD_C; aa 710-848); PF12705 (PDDEXK_1; aa 1072-1214). | Gabija__GajB_2 |
| I55 | *orf25* | RecC | 34334 | 37804 | 3471 | reverse | RecBCD enzyme subunit RecC | PF04257 (Exonuc_V_gamma; aa 11-360); PF17946 (RecC_C; aa 835-1081). | NA |
| I55 | *orf26* | unannotated | 38042 | 38371 | 330 | forward | hypothetical protein | NA | NA |
| I55 | *orf27* | DUF932 | 38462 | 39430 | 969 | forward | hypothetical protein | PF06067 (DUF932; aa 77-302). | NA |
| I55 | *orf28* | YqaJ | 39513 | 40514 | 1002 | forward | hypothetical protein | PF09588 (YqaJ; aa 28-171). | NA |
| I55 | *orf29* | Gp3-like | 40605 | 41498 | 894 | forward | hypothetical protein | PF18897 (Gp3-like; aa 12-206). | NA |
| I55 | *orf30* | unannotated | 41554 | 41742 | 189 | forward | hypothetical protein | NA | NA |
| I55 | *orf31* | unannotated | 41803 | 42126 | 324 | forward | hypothetical protein | NA | NA |
| I55 | *orf32* | HTH-containing protein | 42136 | 42483 | 348 | reverse | hypothetical protein | PF01381 (HTH_3; aa 17-71). | NA |
| I55 | *orf33* | unannotated | 42622 | 42975 | 354 | forward | hypothetical protein | NA | NA |
| I55 | *orf34* | unannotated | 43189 | 43812 | 624 | forward | hypothetical protein | NA | NA |
| I55 | *orf35* | MUG113 | 43980 | 45188 | 1209 | reverse | hypothetical protein | PF13455 (MUG113; aa 285-373). | NA |
| I55 | *orf36* | DEAD-like helicase | 45181 | 47247 | 2067 | reverse | hypothetical protein | PF00270 (DEAD; aa 28-193). | NA |
| I55 | *orf37* | DNA methyltransferase | 47349 | 50105 | 2757 | reverse | hypothetical protein | PF20464 (MmeI_N; aa 1-159); PF20465 (MmeI_hel; aa 170-250); PF20473 (MmeI_Mtase; aa 325-605); PF20466 (MmeI_TRD; aa 627-826); PF20467 (MmeI_C; aa 827-909). | RM_Type_IIG__Type_IIG |

| | | | | | | | | | |
|---|---|---|---|---|---|---|---|---|---|
| I55 | *orf38* | inovirus Gp2 | 50350 | 51321 | 972 | reverse | hypothetical protein | NA | NA |
| I55 | *orf39 (alpA)* | AlpA | 51661 | 51888 | 228 | reverse | hypothetical protein | PF05930 (Phage_AlpA; aa 1-51). | NA |
| I55 | *orf40* | unannotated | 51950 | 53260 | 1311 | reverse | hypothetical protein | NA | NA |
| I55 | *orf41* | unannotated | 53417 | 54289 | 873 | forward | hypothetical protein | NA | NA |

- **ORF name**: This is the name given to a gene on an MGE. On each MGE, genes are named serially and consecutively from 5'-end (near tyrosine integrase) to 3'-end in the format *orf*#, where # is a two-digit number. The tyrosine integrase always has #=01, the genes following it have #=02, 03, 04, etc. For some well-recognisable genes, common gene names are indicated inside round brackets following *orf*#.
- **Consensus annotation**: This is the annotation given to a gene after manual checks on Prokka and Pfam predictions. Genes are called "unannotated" when neither predictors find results.
- **Minimum coordinate**: This is the position of the first nucleotide of a gene. For each MGE, the nucleotides are numbered from the 5'-end, with the first nucleotide immediately downstream of the 14 bp repeat (near the 3'-end of *tmRNA*) labelled as 1.
- **Maximum coordinate**: This is the position of the last nucleotide of a gene.
- **Length**: This is the number of nucleotides in a gene, and is equal to Maximum coordinate - Minimum coordinate + 1.
- **Direction of transcription**: This is the direction of transcription of a gene. Genes labelled as "forward" are transcribed from the 5'-end to the 3'-end along the annotated strand (i.e., the strand where the tyrosine integrase is at the 5'-end). Genes labelled as "forward" are transcribed from the 5'-end to the 3'-end along the other stand.
- **Prokka product**: This is the protein product predicted by Prokka in KBase. Proteins without predictions are labelled as "hypothetical protein".
- **Pfam domains**: This is the protein domain(s) predicted by Pfam in InterProScan. For each predicted domain, the Pfam Id is listed, with the corresponding Pfam name and mapped region (given by the interval of amino acids along the amino acid sequence) given in round brackets. For proteins with multiple predicted domains, all domains are listed and separated by semicolons. Proteins with no predicted domains are labelled as "NA".
- **DefenseFinder prediction**: This is the gene name of putative phage defence genes predicted by DenfenseFinder. Genes not predicted to be involved in phage defence activities are labelled as "NA".

### **Table S5**. Mutations detected in isolates with MGEs.

| Strain number | Genotype | Gene locus | Genome coordinate | Type of mutation | Change | CDS position | Effect on protein |
|---|---|---|---|---|---|---|---|
| MPB38392 | SBW25 *tmRNA*::I23 *glmS*::Tn*7*-152(GFP, Kan$^R$) isolate1 | *pflu4414* | 4878944-4878945 | insertion (tandem repeat) | T(2) -> T(3) | 1489-1490 | frameshift |
| MPB38393 | SBW25 *tmRNA*::I23 *glmS*::Tn*7*-152(GFP, Kan$^R$) isolate2 | *pflu4414 - pflu4415* | 4880381-4880570 | deletion | del 190 bp | *pflu4414*: 1-53; *pflu4415*: 664-789 | *pflu4415*: truncation; *pflu4414*: no translation |
| MPB38394 | SBW25 *tmRNA*::I23 *glmS*::Tn*7*-152(GFP, Kan$^R$) isolate3 | *pflu4414* | 4880259 | SNP (transition) | G -> A | 175 | truncation |
| MPB38395 | SBW25 *tmRNA*::I23 *glmS*::Tn*7*-152(GFP, Kan$^R$) isolate4 | *pflu3022* | 3295100-3295889 | deletion | del 790 bp | 387-1176 | truncation |
| | | *pflu4414* | 4879083-4879084 | insertion (tandem repeat) | T(5) -> T(6) | 1350-1351 | frameshift |
| MPB38396 | SBW25 *tmRNA*::I23 *glmS*::Tn*7*-152(GFP, Kan$^R$) isolate5 | *pflu4414 - pflu4415* | 4880381-4880570 | deletion | del 190 bp | *pflu4414*: 1-53; *pflu4415*: 664-789 | *pflu4415*: truncation; *pflu4414*: no translation |
| MPB38895 | SBW25 *tmRNA*::I23 *glmS*::Tn*7*-155(RFP, Kan$^R$) isolate6 | None | - | - | - | - | - |
| MPB38906 | SBW25 *tmRNA*::I44 *glmS*::Tn*7*-155(RFP, Kan$^R$) isolate1 | *pflu4744* | 5217912 | SNP (transition) | G -> A | 116 | substitution (S39N) |
| MPB38907 | SBW25 *tmRNA*::I44 *glmS*::Tn*7*-155(RFP, Kan$^R$) isolate2 | *pflu4744* | 5217912 | SNP (transition) | G -> A | 116 | substitution (S39N) |
| MPB33357 | SBW25 *tmRNA*::I55 *glmS*::Tn*7*-152(GFP, Kan$^R$) isolate1 | *pflu3805* | 4199776 | SNP (transversion) | A -> T | 33 | substitution (N11K) |
| | | *pflu4414* | 4880277-4880278 | insertion (tandem repeat) | C(5) -> C(6) | 157-158 | frameshift |

**Table S6**. Rank abundance of genes carried by similar MGEs

| Domains | COG category | Description | Counts |
|---|---|---|---|
| Phage_integrase | L | Belongs to the 'phage' integrase family | 111 |
| AlpA | K | transcriptional regulator | 109 |
| Inovirus_Gp2 | S | Inovirus Gp2 | 101 |
| WYL | K | regulation of single-species biofilm formation | 87 |
| DUF932 | S | Domain of unknown function (DUF932) | 80 |
| YqaJ | L | COG5377 Phage-related protein | 78 |
| RadC | L | RadC-like JAB domain | 66 |
| DUF1524,DUF262 | S | Protein of unknown function DUF262 | 42 |
| HsdM_N,N6_Mtase | V | Type I restriction-modification system methyltransferase subunit | 40 |
| Methylase_S | V | PFAM Type I restriction modification DNA specificity domain | 31 |
| N6_Mtase | V | Type II restriction enzyme, methylase | 24 |
| DUF1788 | S | Domain of unknown function (DUF1788) | 23 |
| AAA_5 | V | GTPase subunit of restriction endonuclease | 20 |
| Dynamin_N | T | ATPase. Has a role at an early stage in the morphogenesis of the spore coat | 20 |
| HTH_3,HTH_31 | K | sequence-specific DNA binding | 20 |
| MMR_HSR1 | P | Dynamin family | 17 |
| ADP_ribosyl_GH | O | ADP-ribosylglycohydrolase | 16 |
| DNA_methylase | H | Belongs to the class I-like SAM-binding methyltransferase superfamily. C5-methyltransferase family | 16 |
| DUF262 | S | Protein of unknown function DUF262 | 16 |
| Helicase_C,SNF2_N | KL | Transcription regulator that activates transcription by stimulating RNA polymerase (RNAP) recycling in case of stress conditions such as supercoiled DNA or high salt concentrations. Probably acts by releasing the RNAP, when it is trapped or immobilized on tightly supercoiled DNA. Does not activate transcription on linear DNA. Probably not involved in DNA repair | 15 |
| Vsr | L | May nick specific sequences that contain T G mispairs resulting from m5C-deamination | 15 |
| EcoR124_C,HSDR_N,ResIII | L | Subunit R is required for both nuclease and ATPase activities, but not for modification | 14 |
| Lon_2,Lon_C | O | Putative ATP-dependent Lon protease | 14 |
| PDDEXK_1,UvrD-helicase,UvrD_C | L | A helicase nuclease that prepares dsDNA breaks (DSB) for recombinational DNA repair. Binds to DSBs and unwinds DNA via a highly rapid and processive ATP-dependent bidirectional helicase activity. Unwinds dsDNA until it encounters a Chi (crossover hotspot instigator) sequence from the 3' direction. Cuts ssDNA a few nucleotides 3' to the Chi site. The properties and activities of the enzyme are changed at Chi. The Chi-altered holoenzyme produces a long 3'-ssDNA overhang and facilitates RecA-binding to the ssDNA for homologous DNA recombination and repair. Holoenzyme degrades any linearized DNA that is unable to undergo homologous recombination. In the holoenzyme this subunit contributes ATPase, 3'-5' helicase, exonuclease activity and loads RecA onto ssDNA | 14 |
| Helicase_C | KL | Helicase conserved C-terminal domain | 13 |
| HTH_1,LysR_substrate | K | transcriptional regulator | 13 |

| | | | |
|---|---|---|---|
| McrBC | V | McrBC 5-methylcytosine restriction system component | 13 |
| HSDR_N,ResIII | L | Subunit R is required for both nuclease and ATPase activities, but not for modification | 12 |
| HTH_17 | K | TIGRFAM DNA binding domain, excisionase family | 12 |
| DUF1998 | S | Domain of unknown function (DUF1998) | 11 |
| PglZ | H | PglZ domain | 11 |
| Ank_2,Ank_4,Ank_5 | G | response to abiotic stimulus | 10 |
| N6_N4_Mtase | H | DNA methylase N-4 N-6 | 10 |
| ResIII | KL | Type III restriction enzyme, res subunit | 10 |
| DEAD,DUF1998,Helicase_C | L | COG1205 Distinct helicase family with a unique C-terminal domain including a metal-binding cysteine cluster | 9 |
| DUF1819 | S | Putative inner membrane protein (DUF1819) | 9 |
| DUF45 | S | Metal-dependent hydrolase | 9 |
| HTH_Tnp_1 | L | Transposase and inactivated derivatives | 9 |
| Lon_2,Lon_C,MIT_C | O | ATP-dependent Lon-type protease | 9 |
| PLDc_2 | I | phospholipase d transphosphatidylase | 9 |
| AAA_15 | L | AAA ATPase domain | 8 |
| BPD_transp_1 | P | Binding-protein-dependent transport system inner membrane component | 8 |
| DNA_processg_A | LU | DNA recombination-mediator protein A | 8 |
| DUF697,MMR_HSR1 | D | 50S ribosome-binding GTPase | 8 |
| Transp_cyt_pur | FH | Permease for cytosine/purines, uracil, thiamine, allantoin | 8 |
| AAA_15,AAA_21,AAA_23 | T | AAA domain, putative AbiEii toxin, Type IV TA system | 7 |
| DEAD,Helicase_C,RecQ_Zn_bind | L | DNA helicase | 7 |
| Eco57I,N6_Mtase | V | COG1002 Type II restriction enzyme, methylase subunits | 7 |
| AAA_11,AAA_12,AAA_30,PDDEXK_1,RNase_H_2 | L | RNase_H superfamily | 6 |
| AAA_19,AAA_30,UvrD_C_2 | L | A helicase nuclease that prepares dsDNA breaks (DSB) for recombinational DNA repair. Binds to DSBs and unwinds DNA via a highly rapid and processive ATP-dependent bidirectional helicase activity. Unwinds dsDNA until it encounters a Chi (crossover hotspot instigator) sequence from the 3' direction. Cuts ssDNA a few nucleotides 3' to the Chi site. The properties and activities of the enzyme are changed at Chi. The Chi-altered holoenzyme produces a long 3'-ssDNA overhang and facilitates RecA-binding to the ssDNA for homologous DNA recombination and repair. Holoenzyme degrades any linearized DNA that is unable to undergo homologous recombination. In the holoenzyme this subunit has ssDNA-dependent ATPase and 5'-3' helicase activity. When added to pre-assembled RecBC greatly stimulates nuclease activity and augments holoenzyme processivity. Negatively regulates the RecA-loading ability of RecBCD | 6 |
| AAA_23 | L | ATPase involved in DNA repair | 6 |
| ABC_tran,DUF726,Dynamin_N | S | ATPase. Has a role at an early stage in the morphogenesis of the spore coat | 6 |
| CxxCxxCC | S | PFAM Uncharacterised protein family (UPF0153) | 6 |
| DUF2357,PDDEXK_7 | S | PD-(D/E)XK nuclease superfamily | 6 |

| | | | |
|---|---|---|---|
| DUF445 | S | membrane | 6 |
| EH_Signature | S | EH_Signature domain | 6 |
| Exonuc_V_gamma | L | A helicase nuclease that prepares dsDNA breaks (DSB) for recombinational DNA repair. Binds to DSBs and unwinds DNA via a highly rapid and processive ATP-dependent bidirectional helicase activity. Unwinds dsDNA until it encounters a Chi (crossover hotspot instigator) sequence from the 3' direction. Cuts ssDNA a few nucleotides 3' to the Chi site. The properties and activities of the enzyme are changed at Chi. The Chi-altered holoenzyme produces a long 3'-ssDNA overhang and facilitates RecA-binding to the ssDNA for homologous DNA recombination and repair. Holoenzyme degrades any linearized DNA that is unable to undergo homologous recombination. In the holoenzyme this subunit recognizes the wild- type Chi sequence, and when added to isolated RecB increases its ATP-dependent helicase processivity | 6 |
| Helicase_C,Mrr_cat,SNF2_N, SWIM | KL | COG0553 Superfamily II DNA RNA helicases, SNF2 family | 6 |
| PDDEXK_1 | L | PD-(D/E)XK nuclease superfamily | 6 |
| Amidohydro_3 | F | cytosine deaminase | 5 |
| DDE_Tnp_1,DDE_Tnp_1_6,DUF772 | L | Transposase DDE domain | 5 |
| DUF1232 | S | Protein of unknown function (DUF1232) | 5 |
| DUF3987 | T | Protein of unknown function (DUF3987) | 5 |
| Helicase_C,ResIII,SNF2_N | L | PFAM Helicase conserved C-terminal domain | 5 |
| HNH_2 | V | restriction endonuclease | 5 |
| Hydant_A_N,Hydantoinase_A | EQ | N-methylhydantoinase A acetone carboxylase, beta subunit | 5 |
| Hydantoinase_B | EQ | Hydantoinase B/oxoprolinase | 5 |
| Isochorismatase | Q | Isochorismatase family | 5 |
| MFS_1 | EGP | Major Facilitator Superfamily | 5 |
| MFS_1,Sugar_tr | EGP | Major facilitator superfamily | 5 |
| MotB_plug,OmpA | N | Membrane MotB of proton-channel complex MotA/MotB | 5 |
| PDDEXK_7 | S | PD-(D/E)XK nuclease superfamily | 5 |
| UvrD-helicase,UvrD_C,UvrD_C_2 | L | Uvrd rep helicase | 5 |
| Virulence_RhuM | S | COG3943 Virulence protein | 5 |
| AAA | O | COG0464 ATPases of the AAA class | 4 |
| AAA_13 | L | AAA domain | 4 |
| AAA_5,DUF3578 | V | Domain of unknown function (DUF3578) | 4 |
| ABC_tran,TOBE_2 | P | Belongs to the ABC transporter superfamily | 4 |
| DAO | E | FAD dependent oxidoreductase | 4 |
| DUF2950 | N | Protein of unknown function (DUF2950) | 4 |
| DUF3077 | S | Protein of unknown function (DUF3077) | 4 |
| DUF4357 | S | Domain of unknown function (DUF4357) | 4 |

| | | | |
|---|---|---|---|
| Fer2_4 | C | 2Fe-2S iron-sulfur cluster binding domain | 4 |
| Fer2_BFD,Pyr_redox_2 | C | FAD-dependent pyridine nucleotide-disulphide oxidoreductase | 4 |
| Helicase_C,ResIII | L | Type III restriction protein res subunit | 4 |
| His_Phos_1,NUDIX | L | nUDIX hydrolase | 4 |
| HTH_30,PucR | QT | transcriptional regulator | 4 |
| Ketoacyl-synt_C,ketoacyl-synt | IQ | catalyzes a condensation reaction in fatty acid biosynthesis addition of an acyl acceptor of two carbons from malonyl-ACP | 4 |
| Pkinase | KLT | Protein tyrosine kinase | 4 |
| Ribonuc_L-PSP | J | Endoribonuclease L-PSP | 4 |
| SBP_bac_6,SBP_bac_8 | E | Bacterial extracellular solute-binding protein | 4 |
| AAA_11,AAA_12,NERD,Pkinase | L | AAA domain | 3 |
| Arm-DNA-bind_3,Phage_integrase | L | Belongs to the 'phage' integrase family | 3 |
| DDE_Tnp_IS66,DDE_Tnp_IS66_C,LZ_Tnp_IS66,zf-IS66 | S | zinc-finger binding domain of transposase IS66 | 3 |
| DEAD,Helicase_C | L | helicase superfamily c-terminal domain | 3 |
| DJ-1_PfpI,HTH_18 | K | COG4977 Transcriptional regulator containing an amidase domain and an AraC-type DNA-binding HTH domain | 3 |
| DUF3387,HSDR_N,ResIII | F | Subunit R is required for both nuclease and ATPase activities, but not for modification | 3 |
| Helicase_C,PLDc_2,SNF2_N | L | SNF2 family N-terminal domain | 3 |
| HTH_21,rve | L | Transposase and inactivated derivatives | 3 |
| Mrr_cat | V | Restriction endonuclease | 3 |
| NA37 | S | 37-kD nucleoid-associated bacterial protein | 3 |
| RelA_SpoT | S | Region found in RelA / SpoT proteins | 3 |
| RHH_1 | S | Ribbon-helix-helix protein, copG family | 3 |
| Sel1 | S | Sel1-like repeats. | 3 |
| TnpB_IS66 | L | Transposase | 3 |
| AAA_15,AAA_21,ABC_tran | L | (ABC) transporter | 2 |
| AAA_19,NERD,UvrD_C_2 | L | COG0210 Superfamily I DNA and RNA helicases | 2 |
| AAA_19,UvrD-helicase | L | UvrD/REP helicase N-terminal domain | 2 |
| AAA_19,UvrD-helicase,UvrD_C,UvrD_C_2 | L | UvrD/REP helicase N-terminal domain | 2 |
| Acyl-CoA_dh_1,Acyl-CoA_dh_M,Acyl-CoA_dh_N | I | Acyl-CoA dehydrogenase, middle domain | 2 |
| adh_short_C2 | IQ | PFAM Short-chain dehydrogenase reductase SDR | 2 |
| AIPR | S | AIPR protein | 2 |
| Aldedh | C | Aldehyde dehydrogenase family | 2 |

| | | | |
|---|---|---|---|
| CoA_transf_3 | C | CoA-transferase family III | 2 |
| DUF1837 | S | Domain of unknown function (DUF1837) | 2 |
| DUF3223 | S | Protein of unknown function (DUF3223) | 2 |
| DUF3883 | O | Molecular chaperone. Has ATPase activity | 2 |
| DUF4868 | S | Domain of unknown function (DUF4868) | 2 |
| DUF726 | S | Protein of unknown function (DUF726) | 2 |
| EcoEI_R_C,HSDR_N,Helicase_C,ResIII | L | EcoEI R protein C-terminal | 2 |
| GAF,HATPase_c,HisKA,PAS_2,PHY | T | PAS fold | 2 |
| HATPase_c_3 | L | Histidine kinase-, DNA gyrase B-, and HSP90-like ATPase | 2 |
| HNH | L | HNH nucleases | 2 |
| HpcH_HpaI | G | HpcH/HpaI aldolase/citrate lyase family | 2 |
| HTH_29,rve,rve_2,rve_3 | L | COG2801 Transposase and inactivated derivatives | 2 |
| HTH_31 | K | Helix-turn-helix XRE-family like proteins | 2 |
| HxlR | K | transcriptional regulator | 2 |
| MaoC_dehydrat_N,MaoC_dehydratas | I | N-terminal half of MaoC dehydratase | 2 |
| MUG113,T5orf172 | P | T5orf172 | 2 |
| NmrA | GM | NmrA family | 2 |
| PDDEXK_1,Peptidase_M78,UvrD-helicase,UvrD_C | L | Unwinds DNA duplexes with 3' to 5' polarity with respect to the bound strand and initiates unwinding most effectively when a single-stranded region is present | 2 |
| Plug,TonB_dep_Rec | P | TonB dependent receptor | 2 |
| PRiA4_ORF3 | L | Plasmid pRiA4b ORF-3-like protein | 2 |
| RES | S | RES | 2 |
| Response_reg | T | response regulator | 2 |
| SIR2_2 | F | SIR2-like domain | 2 |
| UvrD_C_2 | L | UvrD-like helicase C-terminal domain | 2 |
| 3HCDH,3HCDH_N | I | 3-hydroxyacyl-coa dehydrogenase | 1 |
| AAA_11,AAA_12,DUF3320,DUF4011,DUF559 | L | AAA domain | 1 |
| AAA_16,GAF,GAF_2,HATPase_c,HisKA,PAS_3,Pkinase | T | Histidine kinase | 1 |
| AAA_18,AAA_33,CPT | S | AAA domain | 1 |
| AAA_19,UvrD-helicase,UvrD_C | L | DNA helicase | 1 |
| AAA_21 | S | AAA domain, putative AbiEii toxin, Type IV TA system | 1 |

| | | | |
|---|---|---|---|
| AAA_30,UvrD_C_2 | L | PIF1-like helicase | 1 |
| Abhydrolase_1,Abhydrolase_6 | I | Alpha beta hydrolase | 1 |
| Abhydrolase_1,Hydrolase_4 | I | hydrolases or acyltransferases (alpha beta hydrolase superfamily) | 1 |
| ABM | S | Antibiotic biosynthesis monooxygenase | 1 |
| AcetylCoA_hyd_C,AcetylCoA_hydro | C | acetyl-CoA hydrolase | 1 |
| Acetyltransf_1,Acetyltransf_10,tRNA_m1G_MT | K | acetyltransferase | 1 |
| ACR_tran | V | TIGRFAM transporter, hydrophobe amphiphile efflux-1 (HAE1) family | 1 |
| Acyl-CoA_dh_1,Acyl-CoA_dh_N | I | acyl-CoA dehydrogenase | 1 |
| Adenylsucc_synt | F | Adenylosuccinate synthetase | 1 |
| ADH_N,ADH_zinc_N | C | Zinc-binding dehydrogenase | 1 |
| ADH_N,ADH_zinc_N,ADH_zinc_N_2 | C | alcohol dehydrogenase | 1 |
| ADH_N_2,ADH_zinc_N | S | N-terminal domain of oxidoreductase | 1 |
| adh_short | S | Belongs to the short-chain dehydrogenases reductases (SDR) family | 1 |
| adh_short,adh_short_C2 | IQ | KR domain | 1 |
| Amidohydro_2 | S | Amidohydrolase | 1 |
| AMP_N,Peptidase_M24 | E | Aminopeptidase P, N-terminal domain | 1 |
| ApbA,ApbA_C | H | Ketopantoate reductase PanE/ApbA C terminal | 1 |
| AraC_binding,HTH_18 | K | Transcriptional regulator | 1 |
| Arginase | E | Belongs to the arginase family | 1 |
| Biotin_lipoyl_2,HlyD_3,HlyD_D23 | M | Belongs to the membrane fusion protein (MFP) (TC 8.A.1) family | 1 |
| CitMHS | C | TIGRFAM citrate H symporter, CitMHS family | 1 |
| Cytochrome_CBB3 | C | photosynthesis | 1 |
| DDE_3,HTH_23,HTH_32,HTH_33 | L | COG3335 Transposase and inactivated derivatives | 1 |
| DDE_Tnp_1 | L | PFAM Protein | 1 |
| DDE_Tnp_1,DUF772 | L | COG3039 Transposase and inactivated derivatives, IS5 family | 1 |
| DDE_Tnp_1,Methylase_S | V | type I restriction modification DNA specificity domain | 1 |
| DEAD,Helicase_C,Pribosyltran,RecQ_Zn_bind,UvrD-helicase,UvrD_C | L | helicase superfamily c-terminal domain | 1 |
| DJ-1_PfpI | S | DJ-1/PfpI family | 1 |

| | | | |
|---|---|---|---|
| DoxX_2 | S | DoxX-like family | 1 |
| DUF1302 | M | Protein of unknown function (DUF1302) | 1 |
| DUF2236 | S | Uncharacterized protein conserved in bacteria (DUF2236) | 1 |
| DUF2867 | S | Protein of unknown function (DUF2867) | 1 |
| DUF2938 | S | Protein of unknown function (DUF2938) | 1 |
| DUF2946 | S | Protein of unknown function (DUF2946) | 1 |
| DUF4113,IMS,IMS_C,IMS_HHH | L | DNA polymerase | 1 |
| DUF4420 | S | Putative PD-(D/E)XK family member, (DUF4420) | 1 |
| DUF928,SIR2_2 | F | 5-carbamoylmethyl uridine residue modification | 1 |
| Dynamin_N,MMR_HSR1,Patatin | M | Esterase of the alpha-beta hydrolase superfamily | 1 |
| EcoEI_R_C,Helicase_C,ResIII | L | EcoEI R protein C-terminal | 1 |
| EcoRII-C | L | EcoRII C terminal | 1 |
| ETF | C | Electron transfer flavoprotein | 1 |
| ETF,ETF_alpha | C | Electron transfer flavoprotein | 1 |
| ETF_QO,FAD_binding_2,NAD_binding_8,Thi4 | C | Electron transfer flavoprotein-ubiquinone oxidoreductase | 1 |
| Fe-ADH | C | Iron-containing alcohol dehydrogenase | 1 |
| Fer4_12,Fer4_14,Radical_SAM | S | Radical SAM superfamily | 1 |
| Fer4_12,Radical_SAM | S | Radical SAM superfamily | 1 |
| Flg_new,LRR_5,LRR_6 | S | regulation of response to stimulus | 1 |
| GAF,GAF_2,GGDEF,PAS_3 | T | GAF PAS GGDEF domain-containing protein | 1 |
| GerE | K | helix_turn_helix, Lux Regulon | 1 |
| GerE,Response_reg,Trans_reg_C | K | response regulator | 1 |
| GntR,UTRA | K | UTRA | 1 |
| Histone_HNS | - | - | 1 |
| HTH_11,WYL | K | HTH domain | 1 |
| HTH_18 | K | AraC Family | 1 |
| HTH_20 | K | transcriptional regulator | 1 |
| HTH_21,rve,rve_3 | L | PFAM Integrase catalytic region | 1 |
| HTH_23,HTH_24,rve | L | Transposase and inactivated derivatives | 1 |
| HTH_26 | K | transcriptional regulator | 1 |
| HTH_3,Peptidase_M78 | K | IrrE N-terminal-like domain | 1 |

| | | | |
|---|---|---|---|
| HTH_IclR,IclR | K | transcriptional regulator | 1 |
| Integrase_1,Phage_int_SAM_2 | L | Belongs to the 'phage' integrase family | 1 |
| Lactamase_B | S | Metallo-beta-lactamase superfamily | 1 |
| Lumazine_bd_2 | S | Putative lumazine-binding | 1 |
| LysR_substrate | K | LysR substrate binding domain | 1 |
| M20_dimer,Peptidase_M20 | S | Peptidase dimerisation domain | 1 |
| MaoC_dehydratas | I | MaoC like domain | 1 |
| MS_channel,cNMP_binding | MT | mechanosensitive ion channel | 1 |
| N6_N4_Mtase,TypeIII_RM_meth | L | DNA methylase | 1 |
| NAD_binding_10,NmrA | GM | NmrA-like family | 1 |
| NERD,zf-C4_Topoisom | L | COG0551 Zn-finger domain associated with topoisomerase type I | 1 |
| NUDIX | L | NUDIX domain | 1 |
| OEP | M | RND efflux system, outer membrane lipoprotein | 1 |
| OmpA | N | Flagellar Motor Protein | 1 |
| OsmC | O | response to oxidative stress | 1 |
| Oxidored_FMN | C | PFAM NADH flavin oxidoreductase NADH oxidase | 1 |
| PCuAC | S | Copper chaperone PCu(A)C | 1 |
| PDDEXK_4 | S | PD-(D/E)XK nuclease superfamily | 1 |
| PKD,Tannase | C | PFAM blue (type 1) copper domain protein | 1 |
| Prok-JAB | S | Prokaryotic homologs of the JAB domain | 1 |
| PrpF | P | protein conserved in bacteria | 1 |
| Radical_SAM | C | Elongator protein 3, MiaB family, Radical SAM | 1 |
| Radical_SAM,TRAM,UPF0004 | J | Catalyzes the methylthiolation of an aspartic acid residue of ribosomal protein S12 | 1 |
| RloB | S | RloB-like protein | 1 |
| RraA-like | H | Catalyzes the aldol cleavage of 4-hydroxy-4-methyl-2- oxoglutarate (HMG) into 2 molecules of pyruvate. Also contains a secondary oxaloacetate (OAA) decarboxylase activity due to the common pyruvate enolate transition state formed following C-C bond cleavage in the retro-aldol and decarboxylation reactions | 1 |
| SCO1-SenC | S | SCO1/SenC | 1 |
| SIR2 | K | NAD-dependent lysine deacetylase and desuccinylase that specifically removes acetyl and succinyl groups on target proteins. Modulates the activities of several proteins which are inactive in their acylated form | 1 |
| TauD | Q | Taurine catabolism dioxygenase TauD, TfdA family | 1 |
| Tautomerase_2 | S | tautomerase | 1 |
| TetR_C_13,TetR_N | K | transcriptional regulator | 1 |
| TetR_N | K | Bacterial regulatory proteins, tetR family | 1 |

| | | | |
|---|---|---|---|
| ThiF | H | ThiF family | 1 |
| Thiolase_C,Thiolase_N | I | Acetyl-CoA acetyltransferase | 1 |
| Thioredoxin | O | OST3 / OST6 family, transporter family | 1 |
| Thymidylate_kin | F | Phosphorylation of dTMP to form dTDP in both de novo and salvage pathways of dTTP synthesis | 1 |
| Transposase_mut | L | Transposase | 1 |
| tRNA_anti-like | S | tRNA_anti-like | 1 |
| UvrD-helicase,UvrD_C | L | UvrD-like helicase C-terminal domain | 1 |
| Z1 | L | Z1 domain | 1 |
